# Stable sound decoding despite modulated sound representation in the auditory cortex

**DOI:** 10.1101/2023.01.31.526457

**Authors:** Akihiro Funamizu, Fred Marbach, Anthony M Zador

## Abstract

The activity of neurons in the auditory cortex is driven by both sounds and non-sensory context. To investigate the neuronal correlates of non-sensory context, we trained head-fixed mice to perform a two-alternative choice auditory task in which either reward or stimulus expectation (prior) was manipulated in blocks. Using two-photon calcium imaging to record populations of single neurons in auditory cortex, we found that both stimulus and reward expectation modulated the activity of these neurons. A linear decoder trained on this population activity could decode stimuli as well or better than predicted by the animal’s performance. Interestingly, the optimal decoder was stable even in the face of variable sensory representations. Neither the context nor the mouse’s choice could be reliably decoded from the recorded neural activity. Our findings suggest that in spite of modulation of auditory cortical activity by task priors, auditory cortex does not represent sufficient information about these priors to exploit them optimally and that decisions in this task require that rapidly changing sensory information be combined with more slowly varying task information extracted and represented in brain regions other than auditory cortex.

## Introduction

Appropriate choices based on sensory stimuli are critical to survival. An animal hears a sound, such as a mouse’s squeak or an owl’s hoot, and must decide whether and how to respond to it. The appropriate response depends not only on what the stimulus is, but also on the behavioral context. This behavioral context includes the animal’s present and previous experience, including its memories about what sounds it has heard and what previous choices were successful. Thus, an animal’s response to sensory stimuli adapts to behavioral context.

Contextual adaptation of neural responses occurs throughout the auditory system, from the cochlea to the auditory cortex and beyond. These adaptations allow for better use of limited resources, such as dynamic range (in the case of feedback to the cochlea) or limited attentional resources^1^. Sound responses in auditory cortex and elsewhere in the auditory stream are also modulated by sound statistics^2^, task engagement^3,4^, movement^5^, spectral attention^6^, and fear^7^. Contextual modulation of sound-evoked responses represents a ubiquitous feature of auditory, as well as non-auditory, sensory representations^8^.

To drive behavior, neural representations formed in the auditory cortex must be “decoded”^9,10^ by the downstream areas to which it projects. Here we address two questions about the decoding of auditory cortical representations. First, we ask whether noise in the cortical representation of auditory stimuli constrained the performance of animals performing an auditory discrimination task. Second, we ask how downstream brain areas can decode neural representations in the auditory cortex if those representations are themselves changing because of contextual adaptation. That is, if a sensory representation in area X changes, how can downstream area Y properly exploit the information from X? Mathematically, this problem arises because the usual formulation in which sensory areas communicate via an “information channel” to downstream areas requires that both the sender and the receiver agree on a (fixed) code, and unilateral changes in that code might be expected to degrade the fidelity of information transfer. Intuitively, one might imagine that changing representations might lead to miscommunication between brain areas, for the same reason that changing the meaning of red and green at traffic lights might disrupt traffic flow.

To address these questions quantitatively, we have developed a two-alternative choice auditory decision-making task in which we could manipulate either of two contextual variables: stimulus probability^11–14^ or reward size^15,16^. To maximize reward in this task, subjects must combine stimulus information with the context: stimulus + context → choice. We operationally define the term “context” in what follows to refer to non-sensory variables, such as stimulus probability or reward amount, that we explicitly measured or controlled during the experiment; this definition excludes the many additional variables (such as the animal’s level of hunger) which we did not measure. Using two-photon calcium imaging to record the simultaneous activity of hundreds of neurons in the auditory cortex while mice were performing either the stimulus-probability or reward-size task, we examined what could be decoded from auditory cortical activity in the face of adapting representations.

Here we report that although changes in both reward and stimulus contexts modulated neural representations of sound in the auditory cortex, the optimal decoder for sound was remarkably invariant to different encodings. In many behavioral sessions, decoding the activity of one or a small handful of neurons matched or exceeded the performance of the animal on a trial-by-trial basis, suggesting that cortical noise did not limit the animals’ performance during this task. By contrast, neither context nor choice could be reliably decoded from auditory cortical activity as behavioral context varied, implying that the animals’ decisions depended on the integration of information represented outside of auditory cortex. Our results demonstrate that sound stimuli are encoded by the auditory cortex, and can be reliably and stably read out by downstream areas, even when the encoding is modulated by behavioral context. The stability of auditory cortical sound decoding suggests that plasticity in brain areas downstream of the auditory cortex likely mediate behavioral adaptation induced by changes in behavioral context.

## Results

We first show that mice exploit changes in behavioral context (reward size or stimulus probability) to optimize choices in an auditory decision task. Then, using two-photon calcium imaging of neuronal activity in primary auditory cortex, we establish that sound-evoked neuronal responses are modulated by changes in behavioral context. Next, we construct decoders of neuronal activity, and show that decoding the activity of a small number of neurons—sometimes even a single neuron—matched or exceeded the performance of the animal. Finally, we show that sound decoding is stable, suggesting that (i) downstream areas of auditory cortex do not require context-dependent change of decoding weights to optimize the sound readout; and (ii) plasticity in downstream areas is essential for context-based reward maximization.

### Mice combine sensory stimulus and context in a perceptual decision-making task

We developed a tone-frequency discrimination task for head-fixed mice (**Figure 1A,B**)^17^. Mice were placed on a cylindrical treadmill facing three lick spouts. To initiate a trial, mice were required to lick the center spout, which triggered the delivery of a “tone cloud” sound stimulus composed of 58 overlapping brief pure tones (each pure tone 0.03 s, total 0.6 s). A 0.5μl water reward was delivered at the center spout at the end of the stimulus, at which point the subject was required to lick the left or right spout depending on whether there were more low (5 – 10 kHz) or high (20 – 40 kHz) frequency tones in the tone cloud. Correct choices were rewarded with a sucrose water reward (5%, 2μl), while incorrect choices resulted in a noise burst (0.2 s). The tone clouds were presented at six levels of difficulty (proportion high tones 0, 0.25, 0.45, 0.55, 0.75, 1).

**Figure 1.**
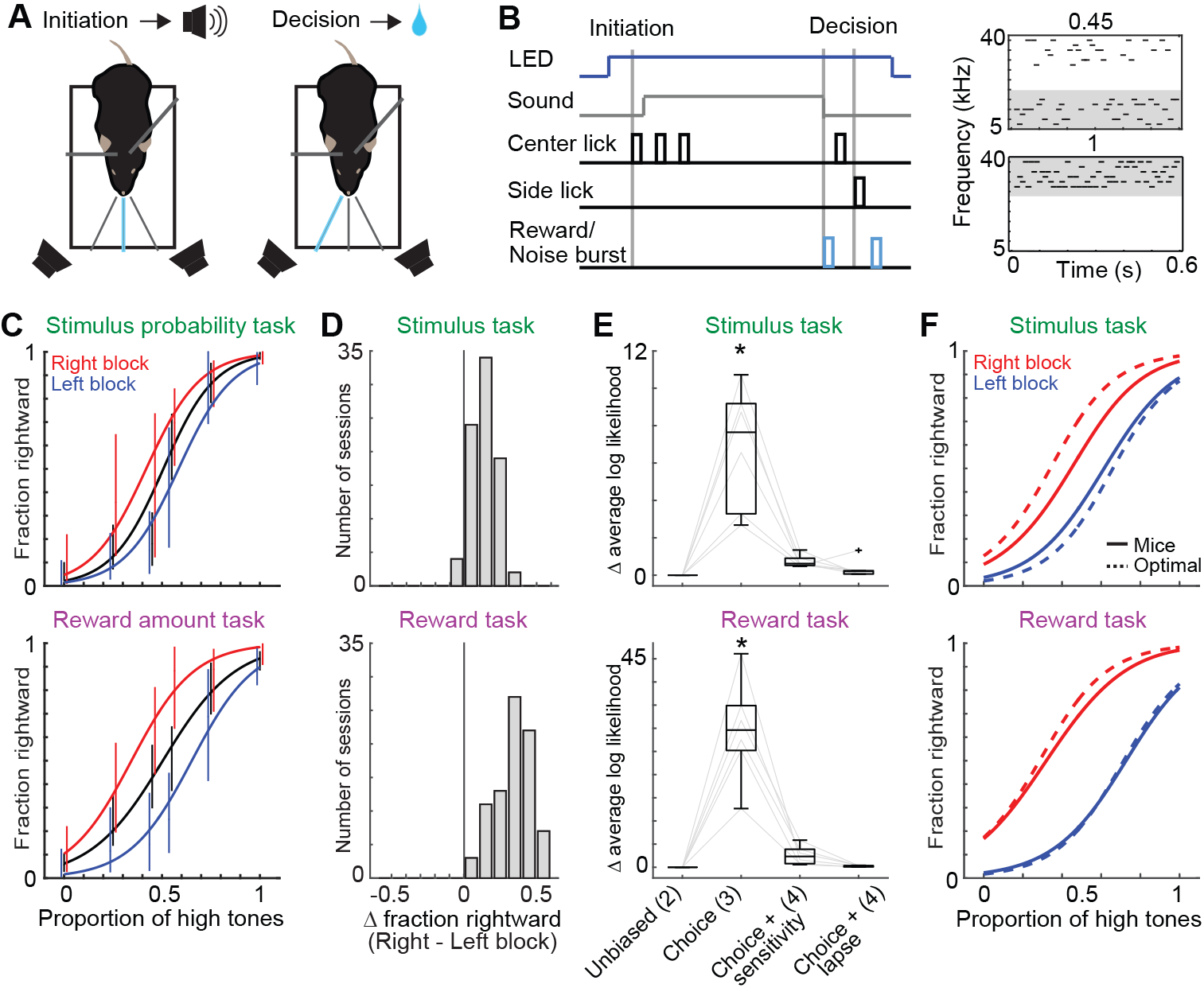
Stimulus probability and reward amount bias choice in auditory discrimination task. (**A**) Behavioral Setup. (**B**) Trial structure. A blue LED indicated the end of the inter-trial interval. Trials were self-initiated: licking the center spout triggered the sound stimulus (0.6 s), at the end of which 0.5 μl of water was delivered on the center spout. Any side-lick thereafter triggered a sucrose reward (correct) or a noise burst (error). Two example stimuli (tone clouds) are shown with a proportion of high frequency tones 45 % (top) and 100 % (bottom). Black lines denote 30 ms pure tones. (**C**) Choice behavior in one session. Error bars show 95 % confidence intervals. The choices were fit with a logistic regression (‘Choice’ in (E)). (**D**) Choice bias in all the 83 sessions. Change in fraction rightward shows the difference of average choice probability in the logistic regression between the left and right blocks (6 mice in each task). (**E**) Choice selective bias. Logistic regression tested whether the choices during the task had no bias depending on blocks (Unbiased), only bias in the choices (Choice), bias in the choices and sound sensitivity (Choice + sensitivity), or bias in the choices with non-zero lapse rate (Choice + lapse). Change in log likelihood in ‘Choice’ showed the increase of log likelihood from ‘Unbiased’. Change in log likelihood in ‘Choice + Sensitivity’ and ‘Choice + Lapse’ showed the increase of log likelihood from ‘Choice’ (83 sessions; central mark in box: median, edge of box: 25th and 75th percentiles, whiskers: most extreme data points not considered outliers (beyond 1.5 times the inter-quartile range), here and hereafter; * p < 0.001 in likelihood ratio test in averaged log likelihood per mouse). Parentheses show the number of parameters in the model. (**F**) Suboptimal choice behavior in stimulus but not reward task. Bold lines show the mean psychometric functions in 83 sessions. Dotted lines show the optimal behavior estimated from the stimulus sensitivity in ‘Choice’ model. See also **Figure S1**.

On alternating sessions, we manipulated either the stimulus probability (the fraction of trials where the stimulus was of the high or low category) or the reward amount. In the stimulus probability task, the stimulus probability for each category alternated between 70%-30% and 30%-70% in blocks of 200 trials (**Figure S1A**). Thus in a 70%-30% block, the stimulus was drawn with 70% probability from one of the three stimuli for which a left response was rewarded (0%, 25%, 45% high-frequency tones). In the reward amount task, the reward amount (3μl or 1μl) associated with correct left and right choices varied in blocks of 200 trials, holding the stimulus probability at 50%-50%. After mice experienced the two asymmetric blocks of stimulus probability or reward amount, these two parameters were set to 50%-50% and 2μl-2μl for the rest of the session.

In each case, the optimal behavior in the face of sensory uncertainty is to make biased decisions. Performance varied smoothly with trial difficulty, with near perfect performance on easy trials (0 or 1 proportion high tones; **Figure 1B,C**) and near chance on difficult trials (0.45 or 0.55). The animal could exploit the context to achieve a higher reward rate especially on the difficult trials. If the animal was in a 30%-70% block where high frequency (rightward) trials are more common than low frequency trials, the mouse should choose rightward more often. Similarly, if a block has more reward on the right (3μl) than left (1μl), the best strategy on difficult trials is to choose rightward. The optimal strategy can be computed based on the task structure (context) and the estimated uncertainty about the stimulus.

We analyzed 83 pairs of stimulus probability and reward amount sessions. Choice behavior was significantly biased to the side associated with the high stimulus probability or large reward amount (p = 1.6E-27 and 5.9E-21 in linear mixed-effects model^18^ in the stimulus and reward tasks, each 83 sessions 6 mice) (**Figure 1D and S1B**), but not by the order of asymmetric blocks (**Figure S1C,D**), confirming that mice used the context information to modify their actions. A logistic regression analysis revealed that the stimulus probability and reward amount affected the choices but did not affect the stimulus sensitivity (slope of psychometric curve) or lapse rate (error rate at 0% and 100% high tones) (**Figure 1E**).

We also assessed whether the mice made optimal use of the context. To compute the optimal strategy, we assumed an ideal observer with a stimulus sensitivity estimated from the mouse’s psychometric curve^19–21^. We found that the subjects’ behavior on the stimulus task was considerably less biased than that predicted from an ideal observer model (linear mixed-effects model, p = 2.3E-25) (**Figure 1F**), but only slightly suboptimal for the reward task (p = 0.091). These are consistent with the observation that the ideal observer obtained more reward than mice (linear mixed-effects model, p = 4.4E-14 and 4.5E-8 in the stimulus and reward tasks, each 83 sessions 6 mice), although mice obtained more reward than an unbiased behavior model with same stimulus sensitivity (p = 0.035 and 1.1E-12). The difference in the observed behavioral bias between the stimulus and reward tasks may arise in part from the fact that subjects can detect a block switch on a single trial for the reward task (because the reward amount on a given port changes by a factor of three), whereas detecting changes in stimulus probability between blocks requires multiple trials. Thus the behavioral data indicate that the mice exploit information about context to increase their reward rate.

### Two-photon microscopy in the auditory cortex during two tasks

We imaged calcium activity from six mice expressing GCaMP6f in excitatory neurons (see Methods) (**Figure 2A**). We used one-photon wide-field imaging to identify the location of primary auditory cortex (**Figure 2B**)^22,23^. In passive animals, 13% (36 / 280) of all neurons per field of view (FOV) showed tone-evoked activity in at least one frequency (**Figure 2C**); the relatively low fraction of tone-responsive neurons is consistent with previous reports^24–26^. We defined a neuron’s best frequency (BF) as the frequency which elicited the highest activity (**Figure 2D and S2A**). To image the activity of the same neurons during both the stimulus and reward tasks, we switched the task on alternate days, keeping the same FOV (**Figure 2E**). We defined “task-relevant neurons” (**Figure 2F**) as any neuron showing increased activity during trial compared to the inter-trial interval in either the stimulus or reward task (see Methods). We identified 13581 task-relevant neurons out of 17523 overlapping neurons (78 % of regions of interest (ROIs) per FOV on average; 6 mice, 83 sessions). The peak activity time and signal strength of task-relevant neurons were correlated across days (**Figure 2G and S2B**)^27^ [for details, see Methods, Supplemental note].

**Figure 2.**
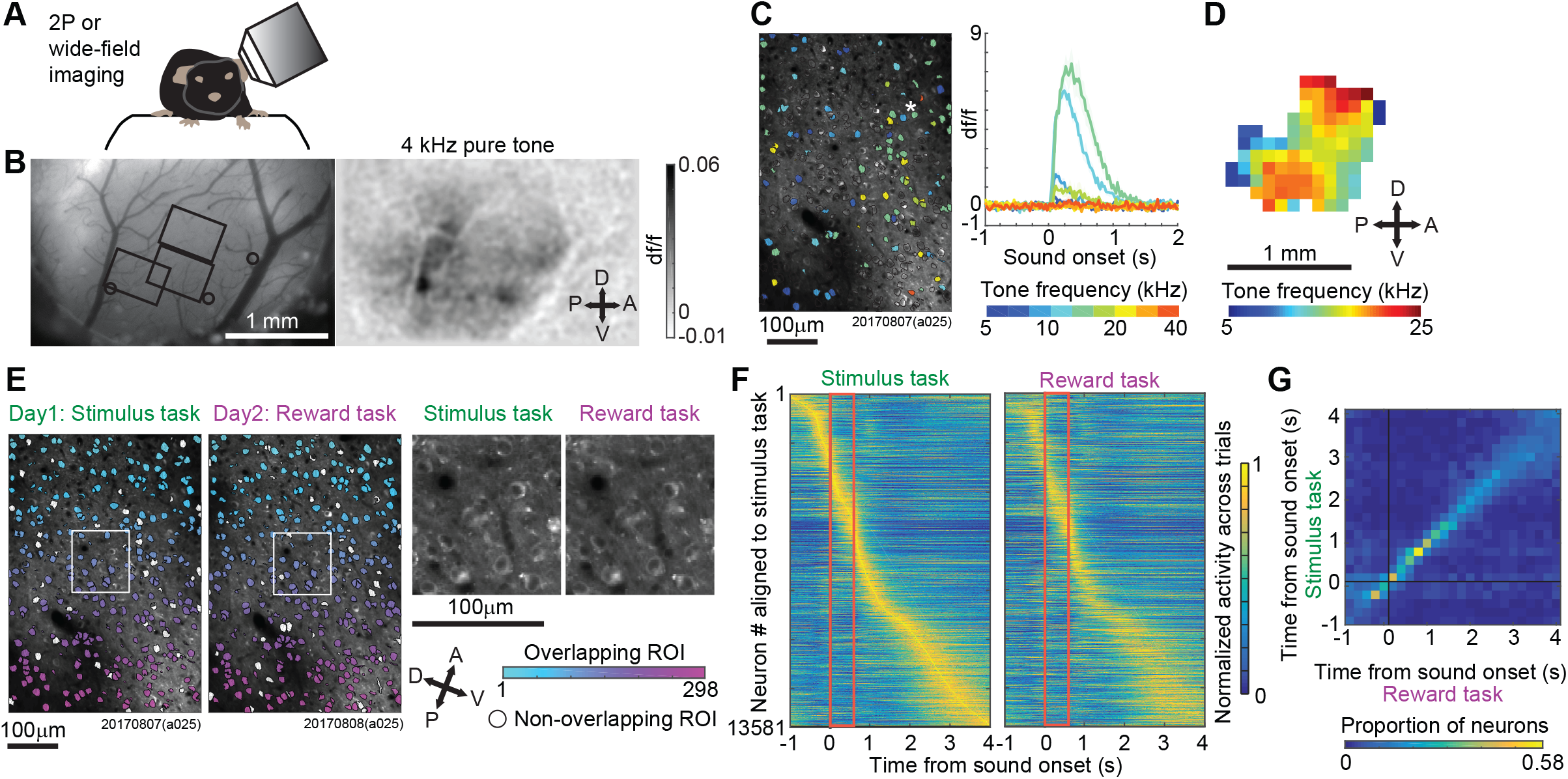
Two-photon microscopy in the auditory cortex during two tasks. (**A**) Setup for microscopy. Objective lens for microscope had a tilted angle to keep the mice parallel to the ground. (**B**) Identification of primary auditory cortex with one-photon wide-field imaging. The 4-kHz pure tone evoked responses through the cranial window provided the locations of the primary, anterior, and secondary auditory fields (right panel). Three circles in the left panel show the approximate peak location of the tone evoked responses. Three rectangles show the imaging locations of two-photon microscopy in the example mouse. The depth of imaging field was different in every session. (**C**) Tone evoked responses in one field of view in two-photon microscopy. The color of each region of interest (ROI) shows the best frequency (BF). ROIs with no color did not show significant sound evoked activity. Example calcium traces with means and standard errors are shown in the ROI with asterisk. (**D**) Sound response map in the imaged location. The BF map of each field of view was merged and superimposed in z axis to show the BF map in xy plane. The BF of each xy location was analyzed as the average BFs of neurons. (**E**) Identification of overlapping ROIs imaged during both the stimulus and reward tasks. The colored and white ROIs were detected as overlapping and non-overlapping, respectively. The magnified view of square site is shown on the right. (**F**) Average activity of task-relevant neurons across trials. The activity was normalized between 0 and 1 and aligned based on the peak activity timings in the stimulus task. (**G**) Peak activity timings of neurons across tasks. Based on the peak timings in reward task (x axis), the proportion of peak timings in stimulus task is shown (y axis; the sum of each column is one) (n = 13581, task-relevant neurons) (Spearman partial correlation eliminating the effect of mice or sessions, time: r = 0.56, p < 1E-10). See also **Figure S2**.

### Sound responses are modulated by the stimulus and reward context

We investigated whether activity in the auditory cortex was modulated by the stimulus or reward context. The median inter-sound-interval was set to 6.6s to minimize task-unrelated sound-evoked activity changes^28^. We analyzed sound-responsive neurons with distinct activity during stimulus presentations and differing activity across tone clouds (**Figure 3A**). 1735 neurons showed sound responses in both tasks. Each neuron’s “preferred tone cloud” was defined as the stimulus eliciting maximum activity. This preference remained consistent between tasks and correlated with passive pure tone presentation frequencies (**Figure S2C,D**). 13% of total neurons responded to sounds, whereas 78% were influenced by the task. Thus, consistent with previous findings in auditory cortex^25^, most task-responsive neurons were not sound-responsive.

**Figure 3.**
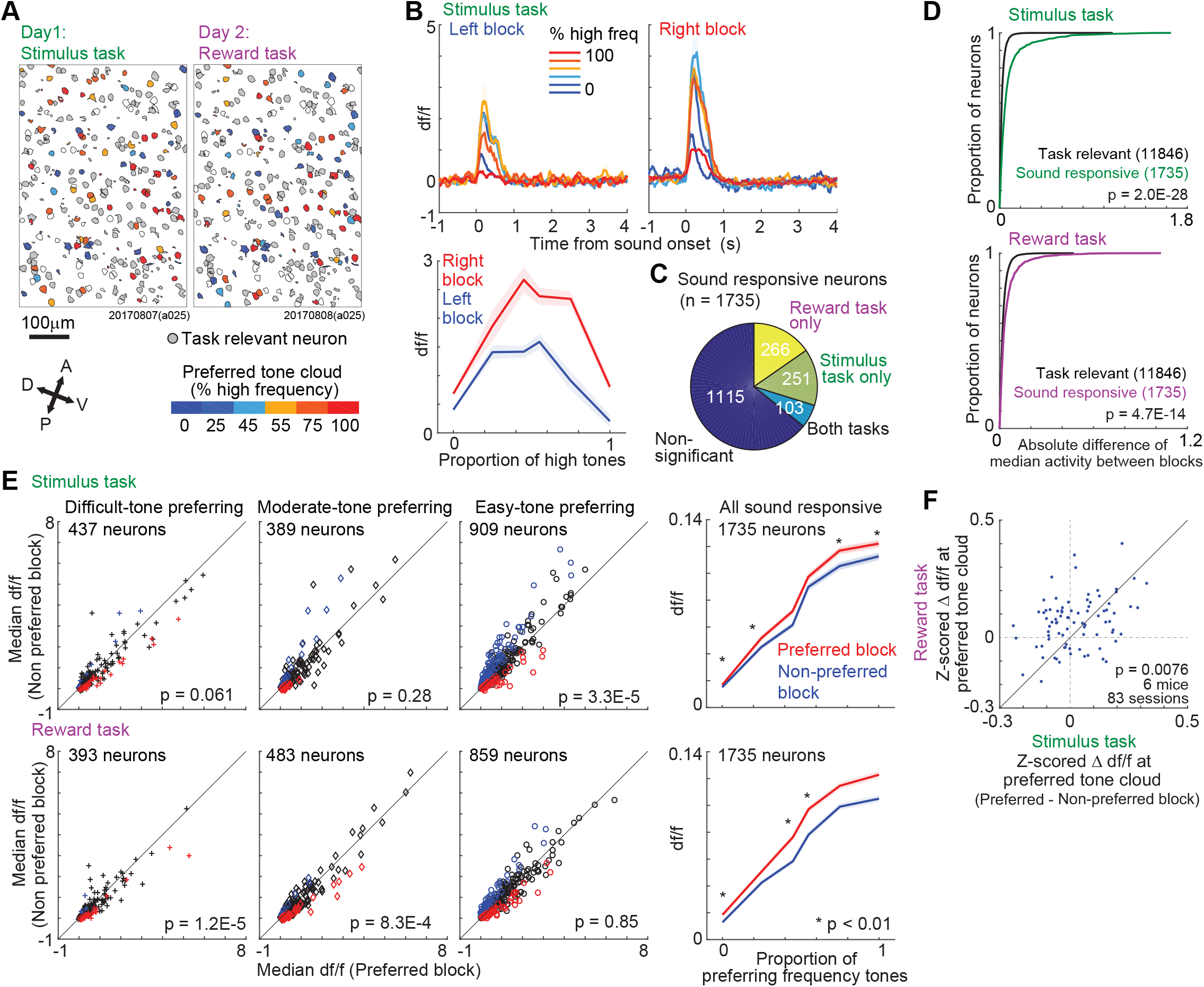
Sound responses are modulated by context about stimulus probability and reward amount. (**A**) Sound responsive neurons in an example field of view (FOV) in Figure 2E. Among the overlapping neurons (white), we defined the task-relevant neurons (gray). Among the task-relevant neurons, we colored neurons according to the preferred tone cloud which elicited the highest activity in sound-responsive neurons. (**B**) Traces of one neuron during left– and right– block trials in the stimulus probability task. The neuron increased the activity in the right block compared to the left block. (Bottom) Tuning curve shows the activity during sounds. Means and standard errors. (**C**) Number of sound responsive neurons with context modulation (p < 0.05, two-sided Mann Whitney U-test at preferred tone cloud). (**D**) Comparison of context modulation of activity between the sound-responsive and task-relevant neurons. Context modulation of sound responsive neurons was greater than sound-unresponsive task-relevant neurons (0.089 +/− 0.16 vs 0.025 +/− 0.030, stimulus modulation; 0.048 +/− 0.075 vs 0.024 +/− 0.027, reward modulation; mean +/− std) (linear mixed-effects model in 6 mice, 83 sessions). (**E**) Context modulation of sound responsive neurons. Sound responsive neurons were categorized to the difficult, moderate, and easy neurons depending on their preferred tone clouds. Scatter plot compared the activity of the preferred tone cloud between blocks. The red and blue points show the significant increase of activity in the preferred and non-preferred block, respectively (p < 0.05 in two-sided Mann-Whitney U test). P-value shows the population comparison with linear mixed-effects model in 6 mice, 83 sessions. The activity was aligned to the preferred tone category. Tuning curve shows the medians and robust standard errors of activity of sound responsive neurons in the preferred and non-preferred blocks. X axis is the proportion of preferred tone category (*, p < 0.01 in linear mixed-effects model). (**F**) Comparison of block-modulated activity between the stimulus and reward tasks. In each session, we analyzed the median block modulation of sound responsive neurons (linear mixed-effects model). See also **Figure S2 and S3**.

We compared the activity between left and right blocks (**Figure 3B**) and found that 35.7% of sound responsive neurons was modulated by the changes in stimulus-probability or reward-amount (**Figure 3C**). Context modulation of sound responsive neurons was greater than sound-unresponsive task-relevant neurons (**Figure 3D**). We then aligned the activity of each sound responsive neuron based on its “preferred block,” i.e. the block associated with the preferred tone cloud stimulus for that neuron. Specifically, the preferred block was “left” for neurons whose preferred tone cloud was in the “low” category and “right” for neurons whose preferred tone cloud was in the “high” category. We categorized the neurons by their preferred tone difficulty and compared activity between blocks elicited during the preferred tone cloud stimulus. We found that the activity could be modulated in either the positive or negative direction (**Figure 3E and S3A,B**), irrespective of choices (**Figure S3C-E**). Modulation was stronger for the reward task (**Figure 3F**). These results indicate that the sound encoding by neurons in the auditory cortex is modulated by stimulus or reward expectation and that the magnitude of the neuronal modulation is correlated with the behavioral bias (**Figure 1D**).

### Neural decoding of stimulus category is comparable to mouse behavior

We next asked how other brain areas could make use of neural representations in auditory cortex to decode the behaviorally relevant stimulus category (low– or high-frequency). We first quantified the decoding performance of single neurons during sound presentation (0.6 s) over the whole session (ignoring the block structure), using a model in which downstream areas decode the stimulus category by setting an optimal threshold on the recorded signal (**Figure 4A**). For each session, the optimal threshold was determined on a training set and then used to classify the neural activity into low and high tones. We found that in a significant fraction of sessions (25% and 55%, respectively, out of 83 sessions in the stimulus and reward tasks), the performance of an ideal observer decoding the best single neuron was better than that of the mouse itself during that session (**Figure 4B-D**), consistent with similar observations in primates^20,29,30^.

**Figure 4.**
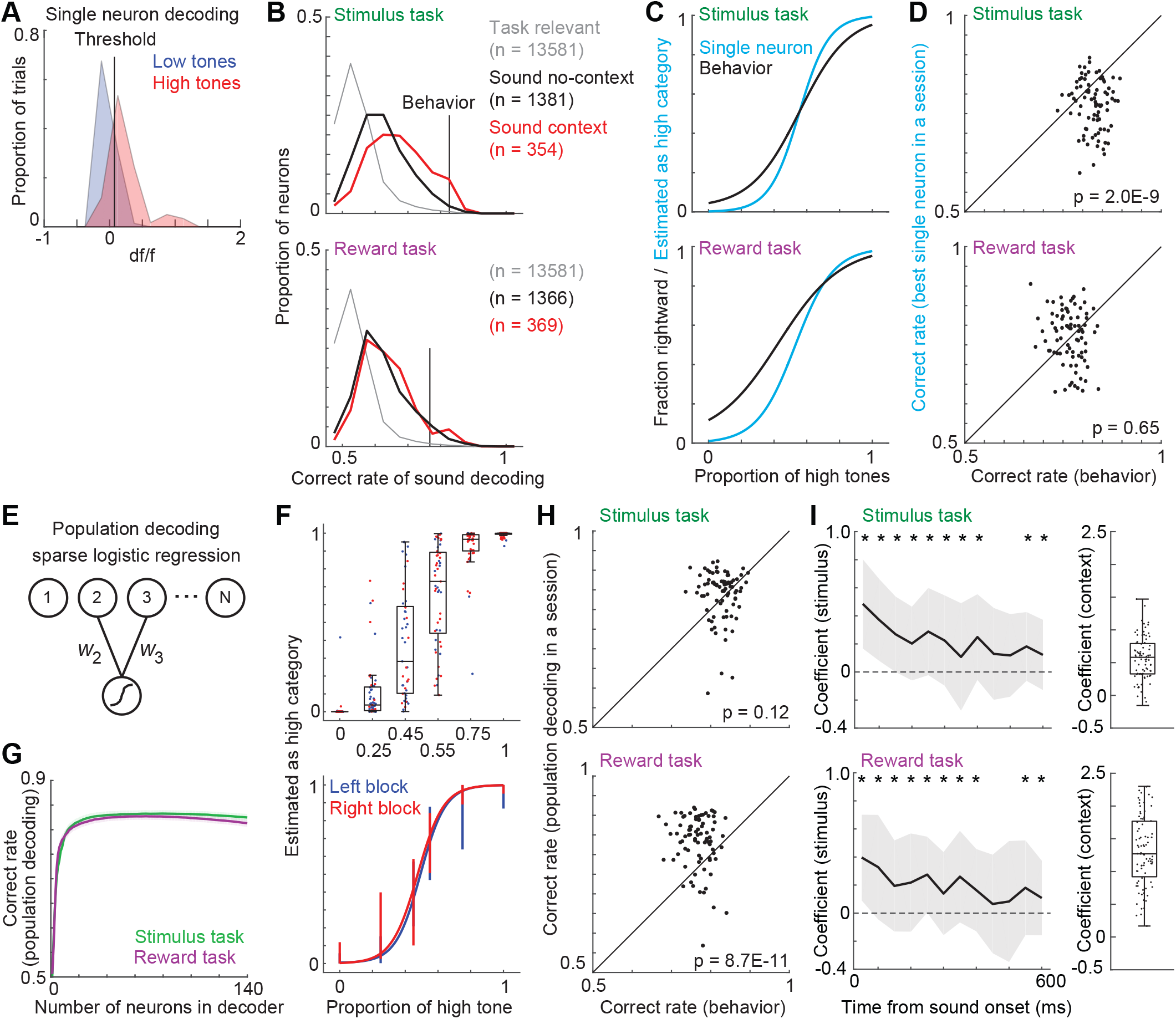
Comparable performance of neural sound decoding and mouse behavior. (**A**) Sound decoding in single neurons. We optimally identified a threshold to discriminate the high– and low-category tones in each neuron (cross validation). (**B**) Distribution of correct rate in single neurons. The vertical bar shows the average correct rate of mice behavior (83 sessions). Sound responsive neurons are separated into context modulated (red) and non-modulated (black). (**C**) Neurometric and psychometric functions in single session. The neurometric function was analyzed from the best sound-decoding neuron in a given session. (**D**) Scatter plot comparing the sound discrimination between the best single neuron and mouse behavior in each session. Single neurons often outperformed the mouse behavior (linear mixed-effects model in 6 mice, 83 sessions). (**E**) Population sound decoding with sparse logistic regression (SLR). SLR sparsely extracted neurons for decoding. (**F**) Sound decoding in one session. Each point shows the estimated probability of high tone category in each trial (400 trials) (top). The probabilistic sound estimation was binarized at 0.5 to analyze the neurometric function. (bottom) Error bars show the 95 % confidence intervals. (**G**) Means and standard errors of correct rate of SLR as a function of number of neurons (83 sessions in each task) (**H**) Comparison of performance between neuronal population and mouse behavior (linear mixed-effects model in 6 mice, 83 sessions). (**I**) Psychophysical kernels. Logistic regression analyzed how tones at each time point and the context information contributed to the choice behavior. Medians and median absolute deviations (left, * p < 0.01 in linear mixed-effects model in 6 mice, 83 sessions). See also **Figure S4**.

We then quantified the decoding performance of the entire neuronal population recorded simultaneously. We used a sparse logistic regression (SLR) decoder, with nested 10-fold cross validation (**Figure 4E**)^31^. We chose the SLR decoder because it performed as well as or better than other decoders tested (**Figure S4A**), and because it has a natural interpretation as a readout by a population of downstream neurons. We again selected the weights optimal for decoding the stimulus category. The neurometric function obtained from this optimal SLR decoder was analyzed by binarizing the probabilistic estimates (**Figure 4F**). To avoid overfitting, the SLR was fitted with an L1 regularizer which identified sparse subsets of neurons for decoding (**Figure 4G**). Population decoding of sound was better on correct trials than on error trials (**Figure S4B**).

As expected, decoding by the population was better than decoding by single neurons (**Figure S4C**), indicating that the sound representation was distributed across the population. Population decoding was often better than the performance of the animal, even when the biases were not exploited by the neural decoder (**Figure 4H)**. That is, the decoder was trained and tested using all trials, regardless of whether they were from a left or right block. All the decoding performance was tested using neuronal activity elicited during sound presentation (0.6 s).

One possible concern is that this relatively long time window might exceed the window over which mice accumulate sound evidence and thus might provide the optimal neural decoder with an unrealistic advantage compared with the mouse’s behavior (**Figure S4D-G**). However, behavioral analysis suggested that mice did accumulate evidence over the entire sound presentation (**Figure 4I**), suggesting that the accumulation was not responsible for the superior performance of the neural decoder. Thus, the optimal readout of both single neurons and neural populations often matched or exceeded the mouse’s performance on a given session.

### Noise correlation among neurons with similar tuning constrains sound decoding performance

Although as expected population decoding was consistently better than single-neuron decoding on single trials (**Figure S4C**), in some cases the advantage gained from using extra neurons in the decoder was relatively small. If extra neurons each carried independent information about the stimulus, then we would expect the inclusion of additional neurons to improve the performance of the decoder; but if the information represented by different neurons were redundant, then the decoding performance would saturate. We therefore examined the role of single-trial correlations in limiting the readout of the stimulus from the neuronal population^32–34^.

We first investigated the “signal correlations”, defined as the correlation between the average stimulus-evoked calcium activity for each of the 6 tone clouds, for all sound-responsive neurons. As expected, signal correlations were higher among neurons with the same preferred tone cloud than neurons with different preferences (example session **Figure 5A**; 83 sessions **Figure 5B**). We then explored noise correlations and found a similar tendency. The activity of neurons with similar stimulus-driven responses were more correlated on single trials than those with different responses (**Figure 5A,C**). Noise correlations were higher between neurons with same preferred stimulus independent of the distance between them (linear mixed-effects model, p = 0.53 – 8.4E-16), indicating the higher noise correlation was not simply a consequence of the tendency of similarly tuned neurons to be closer on the tonotopic map.

**Figure 5.**
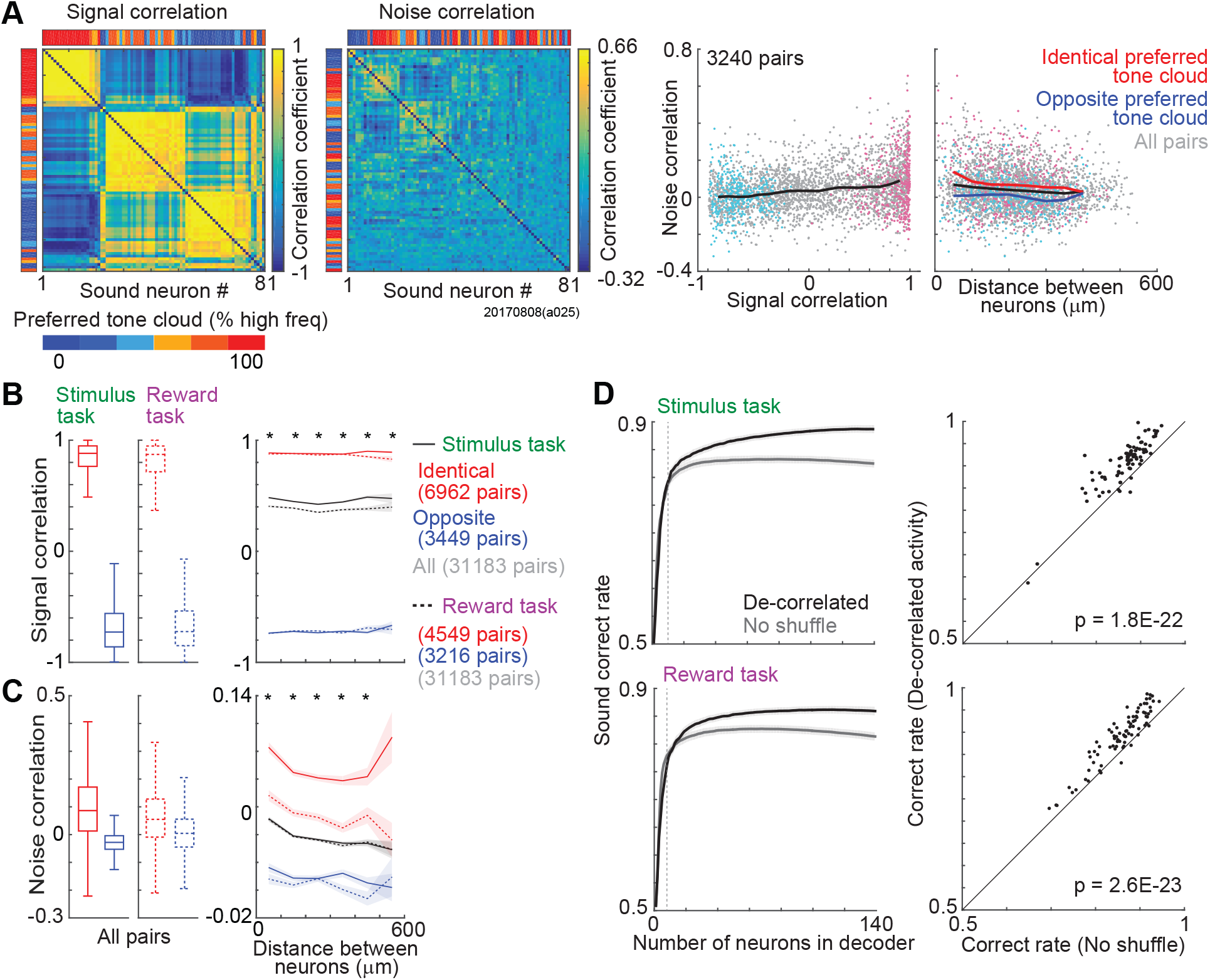
Noise correlations limit sound decoding. (**A**) Signal and noise correlations in one session. Cross correlogram shows the correlations between sound responsive neurons, sorted using hierarchical clustering. Color bars at the top and left show the preferred tone cloud of neurons. Scatter plot shows the relationships among signal correlations, noise correlations, and the physical distance between neurons. Red and blue dots show the data from neurons with same and different preferred tone clouds, respectively (easy stimuli only). Lines show the moving average. (**B**) Signal correlation in 83 sessions. (Left) Box plots compared the signal correlation between neuron pairs of same and different preferred tone clouds (p < 1E-10 in linear mixed-effects model in 6 mice, 83 sessions). (Right) Signal correlation in all neuron pairs as a function of distance between neurons (binned every 100μm of distance between neurons) (* p < 0.001 in linear mixed-effects model in 6 mice, 83 sessions). (**C**) Noise correlations, as in **B**. High noise correlations in the neuron pairs with same preferred tone cloud were observed irrespective of distances (Left: p = 1.3E-25 and 2.1E-18 in linear mixed-effects model in 6 mice, 83 sessions; Right, * p < 0.001). (**D**) Noise correlations limit performance. Means and standard errors (left). Vertical dot line shows the median number of neurons achieved the 95% of maximum correct rate of population decoding. We extracted the neurons achieved the highest correct rate in non-shuffled activity and compared the correct rate with the de-correlated activity (right) (linear mixed-effects model in 6 mice, 83 sessions).

To assess the effect of these noise correlations on stimulus decoding, we compared the performance of a decoder acting on single-trial activity to the performance of a decoder with access to uncorrelated activity obtained on scrambled trials. As expected, decoding based on scrambled trials continued to improve as more neurons were included in the decoder, whereas the performance of the single trial decoder reached an asymptote after around 10-20 neurons and reached 95% of maximum correct rate with 8 neurons on median (**Figure 5D**). These results confirm the role of noise correlations in limiting the potential for reading out activity from a population of neurons. However, the fact that the performance of even the single-trial population decoder often outperformed that of the animal itself (**Figure 4**) suggests that these noise correlations were often not the sole or even main factor in limiting the animals’ performance.

### Optimal decoder for stimulus category is stable

Our previous analyses indicate that cortical noise is not the sole or even main factor limiting the performance of the animal on this task, suggesting that the performance is limited by representations outside of auditory cortex. This raises the question: How can downstream brain areas decode neural representations in the auditory cortex, if those representations are themselves changing because of contextual adaptation? One might imagine that changing representations might lead to miscommunication between brain areas, for the same reason that changing the meaning of red and green at traffic lights might disrupt the flow of traffic.

To maximize reward in this task, an ideal observer (as a model of areas downstream of auditory cortex) with access to the neural activity in the auditory cortex, and with perfect knowledge of context, would make choices by combining auditory activity and stimulus context as follows:

*Optimal_choice* = ***F*** [*Cortical_sound_representation*(stimulus, context), context].

In this equation, the optimal choice is some function ***F*** of both the population response and the context. Context enters into the formation of optimal choice in two ways. First, it enters implicitly, by changing the neural representation of the sound itself, via the term “*Cortical_sound_representation*” (**Figure 3**). Second, context enters explicitly, by changing the optimal action for a given best estimate of sound category. For example, if the neural response encoding the auditory stimulus is completely ambiguous, then a context in which the higher reward is at the left port would dictate that the optimal choice would be “left”. Thus, the optimal choice can be formed by first estimating the stimulus category from the neural response (given the context), and then selecting the choice that maximizes the reward (given the estimate of the stimulus category). Below we consider these two potential effects of context separately.

We first consider the *implicit* effect of context, through its action on the neural encoding of the stimulus. In principle, the optimal strategy for decoding the stimulus category from the context-modulated neural response would be to use different decoders for each of the two contexts. To quantify the effect of context on decoding, we therefore compared the performance of a “dynamic weights” decoding strategy to that of an invariant “constant weights” decoder (**Figure 6A**). When an invariant “constant weights” decoder (trained on data from both blocks) was used, performance was similar across blocks (**Figure 6B**; linear mixed-effects model in 6 mice, 83 sessions, p = 0.073 and 0.079 in the stimulus and reward tasks). Moreover, sound decoding did not improve with a “dynamic weights” decoder in which weights were trained and tested with data within a block (**Figure 6C**; in this as in all analyses, performance was tested on out-of-sample trials, i.e. samples not used for training). Even using different blocks for training and testing (e.g. trained with trials from the left block, and tested with trials from the right block) led to only a modest decline in performance compared with the dynamic weights decoder (**Figure 6D**; 1.6% and 0.37% on median in the stimulus and reward tasks). Furthermore, the stimulus category was identified more accurately than the correct choice (**Figure S5A,B**). Choice decoding reached a maximum after delivery of the reward or noise burst (**Figure S6**) (see Methods). These analyses indicate that, despite changes in sound-evoked responses induced by manipulating context, the identity of the sound can be read out effectively by an invariant decoder that does not adapt to these manipulations.

**Figure 6.**
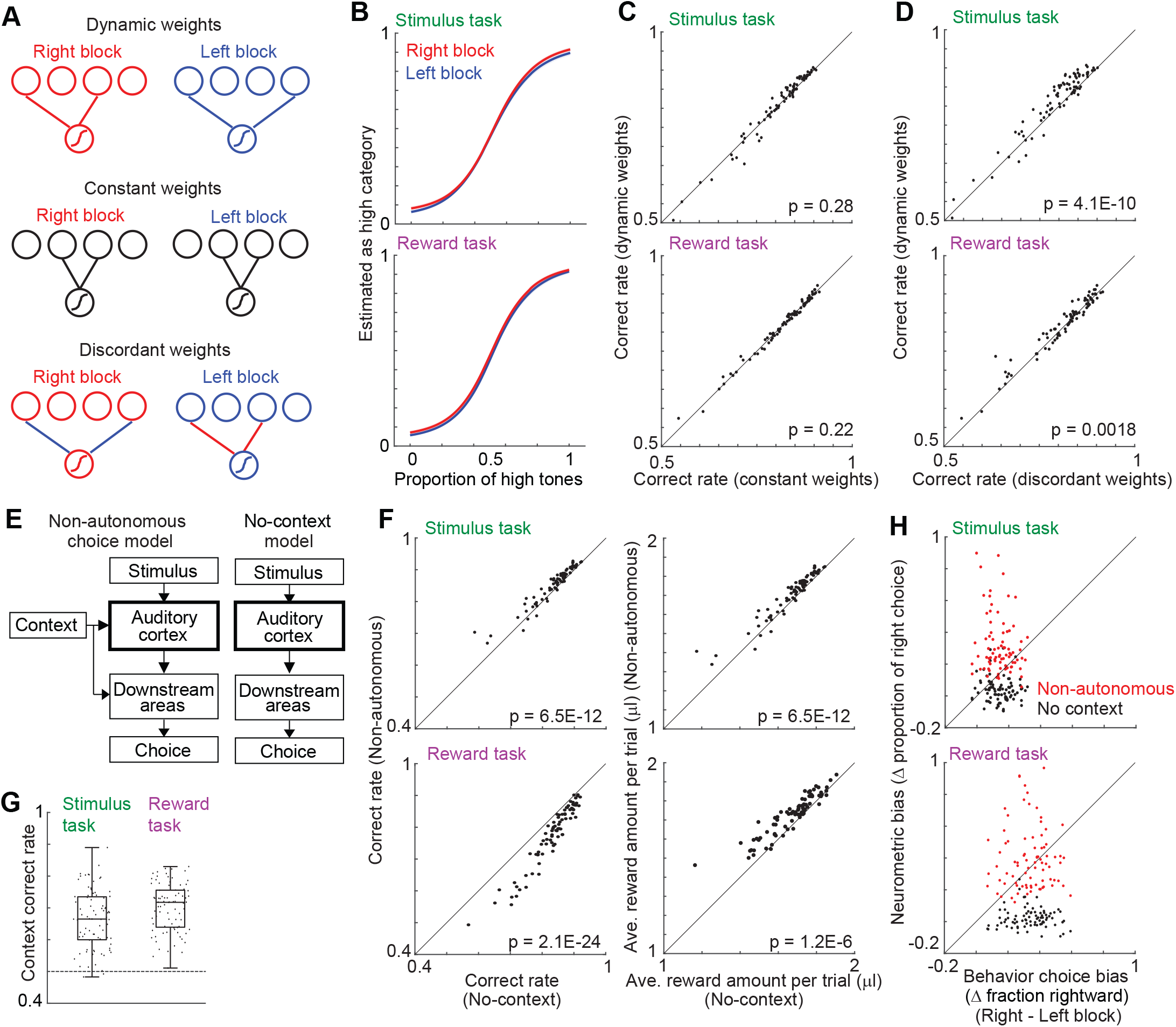
Stable sound readout from the auditory cortex. (**A**) Scheme of sound decoders. Decoder with dynamic weights trained and tested in the same block (optimal decoder). Constant weights had one series of weights across blocks. Discordant weights trained and tested with different blocks (e.g., trained the decoder in left block and tested in right block). (**B**) Means and standard errors of neuromeric functions in constant decoder (6 mice, 83 sessions in each task). (**C, D**) Comparison of decoding performance (6 mice, 83 sessions). Sound decoding with constant weights had comparable performance with the dynamic weights (linear mixed-effects model). (**E**) Two models for how the mouse exploits context for choice. The non-autonomous choice model and no-context model assumed the perfect and no knowledge of context, respectively. (**F**) Comparison of decoding performance in correct rate of sound category (left) and received reward amount (right) (linear mixed-effects model in 6 mice, 83 sessions) (**G**) Context decoding from the auditory cortex (6 mice, 83 sessions). (**H**) Biases in the neuromeric functions across blocks compared with those in the mice behaviors. The bias was the difference of average right choice probability in neuromeric functions between blocks (linear mixed-effects model in 6 mice, 83 sessions: non-autonomous model, p = 1.7E-4 and 0.086 in the stimulus and reward tasks; no-context model, p = 5.6E-6 and 2.3E-17). See also **Figure S5 and S6**.

We next consider the *explicit* effect of context on choice. We first compared the performance of a decoding model which can exploit perfect knowledge of the context (“non-autonomous choice model”) with one that does not exploit context (“no-context model”) (**Figure 6E,F**). Both models first decoded the tone category from the activity in the auditory cortex. The non-autonomous choice model used the optimal decision threshold to compute choices, while the no-context model directly used the outputs of sound decoding as the choices. This comparison revealed the large impact that context can have on this task. The non-autonomous choice model achieved a larger amount of reward compared to the no-context model in 74 and 78 out of 83 sessions in the stimulus and reward task (**Figure 6F** *right*). However, since making an optimal choice in this task requires perfect knowledge of the context, we next attempted to determine the context using only information available from the auditory cortex (“autonomous choice model”). Using the SLR decoder with 10-fold cross validation, we found that the context could only be imperfectly decoded from the population activity (**Figure 6G**, 66% and 70% on average, chance level 50%). In general, stimulus decoding was consistently better than context decoding (**Figure S5C**), and the context decoded from the auditory cortex was insufficient to account for the observed bias in the neurometric function across blocks (**Figure S5D**). In other words, it does not appear that the auditory cortex represents the behavioral context well enough to account for the observed context-dependent shifts in the psychometric curves. Taken together with the invariance of sound decoding, these analyses suggest that the mouse makes choices by combining the auditory stimulus with a representation about context which is encoded downstream (or outside) of the auditory cortex. We compared the shifts in the psychometric curve predicted by the non-autonomous choice model and no-context model to the observed behavior of the mice (**Figure 6H**). For both tasks, the observed behavior was intermediate between the two models. This suggests that, consistent with **Figure 1F**, the animal makes suboptimal use of the block structure of the task, but exploits more of the available context information than is easily decoded from activity in auditory cortex.

### Shifts of decoding threshold are small compared to contextual modulation of neurons

The stability of the stimulus *decoder*, given the variability of the neural *encoding*, seems to pose a conundrum since we might expect that the optimal decoder would vary as the sound representation was modulated by changes in context. One straightforward resolution of this conundrum would be if the decoder relied only on neurons whose activity was not modulated by context. However, although as expected sound decoding relied mainly on sound-responsive neurons (**Figure S7A**), the decoder relied on both neurons modulated by context and those not modulated by context (**Figure 7A**). Thus, stable sound decoding was not achieved by relying only on neurons not modulated by context.

**Figure 7.**
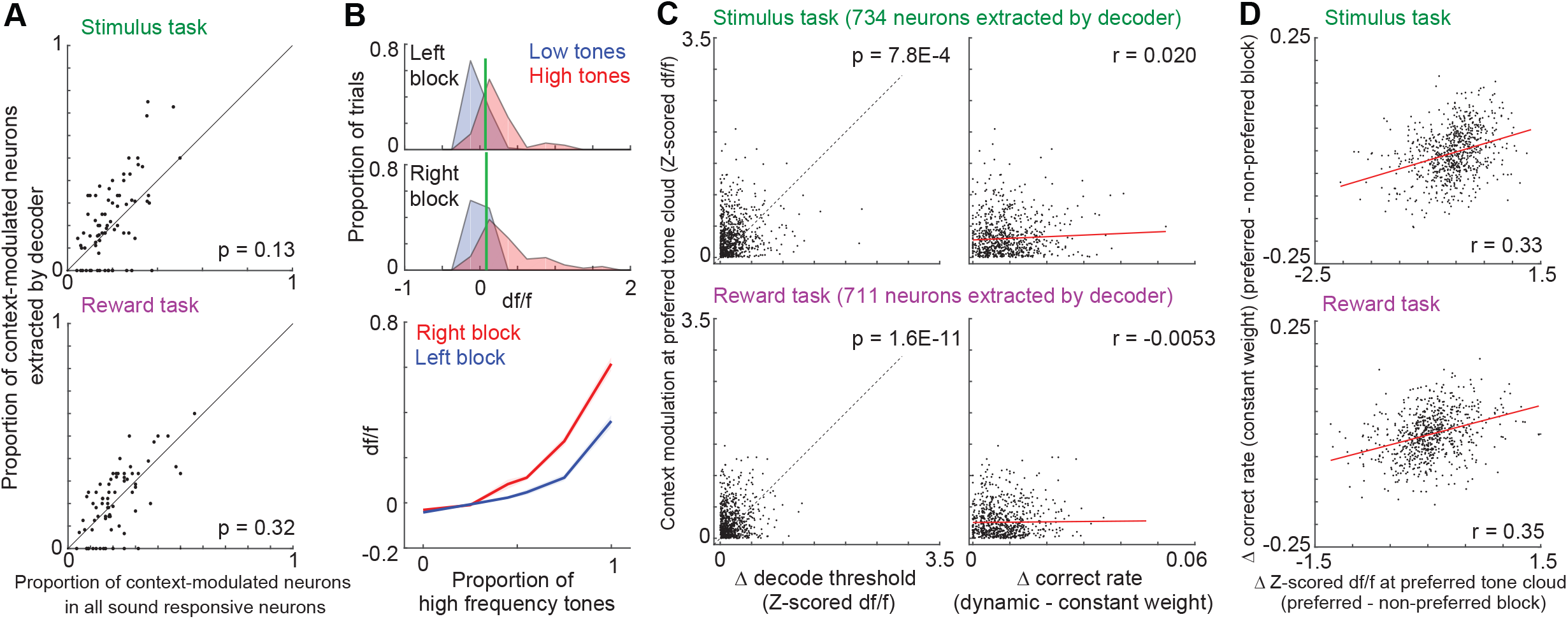
Context modulation of sound responses does not disrupt stable sound decoding. (**A**) Proportion of context-modulated sound responsive neurons extracted in sparse logistic regression (SLR) compared with that in all sound responsive neurons (linear mixed-effects model in 6 mice, 83 sessions in each task). (**B**) Representative neuron with context-modulated activity and stable decoding filter. (top) Decoding thresholds (green lines) which separated the distribution of activity during low and high tones were similar between blocks. (bottom) Contextual modulation of activity. Data presentation same as in Figure 3B. (**C**) Decoding threshold, contextual modulation, and correct rate of sound responsive neurons extracted in the decoder. The contextual modulations were larger than the change in decoding threshold between blocks (left) (linear mixed-effects model in 6 mice, 83 sessions). Correlation between the decoding performance and contextual modulation (right) (Spearman partial correlation, p = 0.60 and 0.89). Contextual modulation was the absolute difference of neural activity at preferred tone cloud between blocks. (**D**) Contextual modulation improved decoding performance. Δ correct rate and Δ Z-scored df/f show the difference of correct rate and the difference of activity between preferred and non-preferred block, respectively (Spearman partial correlation, p = 4.5E-20 and p = 1.6E-21 in the stimulus and reward tasks). See also **Figure S7**.

To resolve this apparent conundrum, we investigated the relationship between sound encoding and decoding in single neurons. **Figure 7B** shows a representative neuron with strong stimulus contextual modulation. However, even though contextual modulation was strong (**Figure 7B** *bottom*), the decoding threshold (*vertical green line, df/f*) was almost unchanged, suggesting that the change in decoding thresholds was small compared to the contextual modulation. This relationship was observed across the population of sound-responsive neurons extracted in the decoder **(Figure 7C** *left* and **S7B**), because the contextual modulations and changes in decoding thresholds were weakly correlated (Spearman partial correlation, r = 0.13 and 0.070 in the two tasks). The contextual modulations did not improve the decoding performance with dynamic weights (**Figure 7C** *right*). In these analyses, the decoding performance was estimated without cross validation to verify that the decoding with block-dependent weights was always better than with a constant weight. These results indicate that the decoding filter of each neuron was relatively stable compared to their contextual modulation. One possible role of the contextual modulation with the stable decoding filter was to improve the decoding performance (**Figure 7D** and **S7C**).

## Discussion

We have used two-photon calcium imaging to record the simultaneous activity of hundreds of neurons in auditory cortex in mice performing a context-dependent two-alternative choice auditory decision task. We find that (1) both the animal’s behavior and neuronal activity are context-dependent; (2) the activity of single neurons in auditory cortex can often be decoded to yield performance as good or even better than the animal, and adding additional neurons often leads to relatively minor improvements in performance; (3) the optimal stimulus decoder remains largely invariant, in spite of context-dependent changes in encoding; and (4) the context decoded from the auditory cortex was insufficient to account for the animal’s behavior. Our results suggest a model in which downstream areas can easily read out information about the stimulus from the activity in auditory cortex, in spite of context-dependent changes in activity.

Since the earliest recordings in auditory cortex, it has been clear that neuronal activity in auditory cortex is strongly modulated by non-sensory features. Hubel described neurons that “appear to be sensitive to auditory stimuli only if the cat ‘pays attention’ to the sound source”^1^. Subsequent studies revealed that neural responses are modulated by sound statistics, attention, task engagement and reward expectation^1–3,5–7,35^. Reinforcing the importance of such contextual modulation, we found that only 13% of neurons responded to tones presented during passive listening, whereas 78% of neurons responded to some component of the task, and about a third (36%) of sound-modulated neurons were modulated by changes in either reward amount or stimulus probability. This modulation raised questions about how downstream brain areas could reliably decode stimulus identity.

To address these issues, we adopted a decoding approach^36^, and assessed how well an ideal observer, with access to activity of hundreds of auditory cortex neurons, could perform on this auditory decision task. In pioneering experiments, Newsome and colleagues related the activity of pairs of neurons in monkey area MT to decisions about motion direction^29,30^. We found, in agreement with these early results, that single neurons could be decoded to yield performance comparable to that of the animal^20,37,38^. This raised the question of why decoding the activity of multiple neurons simultaneously would not do even better. Newsome and colleagues, extrapolating from pairs of neurons, concluded that correlations among neurons limited decoding fidelity. In principle, such correlations could increase or decrease decodability of a population compared with uncorrelated activity, depending on the nature of the correlations^34,39^. Recent recordings of large neuronal populations with two-photon imaging have extended these results beyond pairs of neurons in the context of stimulus encoding^40^. Here we have found that the same principles apply in auditory cortex: We have shown directly, in behaving animals, that decoding the activity of hundreds of auditory neurons simultaneously does not dramatically increase the neurometric performance, compared with decoding the activity of one or just a few of the best neurons (**Figure 5D**).

Given the substantial fraction of neurons modulated by context in this task^8,41–46^, we expected that the optimal decoding filter would vary in order to adapt to this modulation. Surprisingly, however, we found that a single linear decoder performed as well as one that adapted from block to block; the representation of stimulus was orthogonal to the representation of context^47^. In neural terms, this implies that there is no need for hypothetical downstream brain areas decoding the stimulus using the output of auditory cortex to “know” the behavioral context. On the other hand, at the behavioral level mice do exploit behavioral context in this task to maximize reward (**Figure 1**). This implies that the behavioral context is combined with stimulus information outside of the primary auditory cortex (**Figure 6**). Candidate brain areas are the medial prefrontal cortex^48^, parietal cortex^11,49,50^, retrosplenial cortex^51^, and anterior striatum^52^, and secondary auditory cortex^4,53–57^, where neurons have been shown to be modulated by stimulus or reward expectation. Our study also points to the need for exploring high-resolution methodologies like high-density silicon probes^58,59^ to bridge the gap in temporal resolution compared to two-photon calcium imaging (**Figure S4D-G**). Although definitive determination of the causal role of the auditory cortex and downstream regions in perceptual decisions requires manipulations of activity, the decodability of the representations in the face of modulation observed in this study was nonetheless suggestive.

The fact that a simple context-invariant linear model can effectively decode stimuli in the face of context-dependent modulation of activity provides clues as to how the cortex represents stimuli and context. Further experiments are needed to determine whether our results generalize to other sources of contextual modulations such as thirst, attention, and task engagement^1,3,5–7,59^. It would also be of interest to explore how contextual modulation of auditory neurons is acquired by imaging before, during, and after training.

## Acknowledgements

We thank S. Musall for comments, B. Burbach for technical assistance, J. Cohen and V.R. Aguillon for behavior training, H. Zeng for providing Ai93 mice. This work was supported by Uehara Foundation (A.F.), the Watson School of Biological Sciences Farish-Gehry Fellowship (F.M.) and the National Institutes of Health (R01NS088649) (A.Z).

## Author contributions

A.F., F.M. and A.M.Z. designed the experiments. F.M. built the setup. A.F. and F.M. performed the experiments. A.F. analyzed the data. A.F., A.Z. wrote the paper, with comments by F.M.

## Inclusion and diversity

We support inclusive, diverse, and equitable conduct of research.

## STAR Methods

### Resource availability

#### Lead contact

Further information and requests for resources should be directed to Akihiro Funamizu.

#### Materials availability

This study did not generate new materials.

#### Data and code availability

Data analyses were conducted in Matlab scripts. Original codes are going to be publicly available as of the date of publication on Github. The link is listed in the key resources table. The imaging data (df/f) are deposited in Zenodo. The link is in the key resources table. The other imaging data which are not deposited because of the large data size (> 1 TB) are available from the lead contact upon request.

### Experimental model and study participant details

All animal procedures were approved by the Cold Spring Harbor Laboratory Animal Care and Use Committee in an Association for Assessment and Accreditation of Laboratory Animal Care (AAALAC International)-accredited facility and carried out in accordance with National Institutes of Health standards. Mice were housed in a temperature-controlled room with non-inverted, normal 12h/12h light/dark cycle.

#### Chronic window preparation

We used 6 male transgenic GCaMP6f mice (ai93+/–; lsl-tTA+/+; emx-cre+/–) (ai93, Jax stock 024103; lsl-tTA, Jax stock 008600; emx-cre, Jax stock 005628), 8 to 20 weeks of age ^60–63^. Before surgery, mice were restricted to 1.5 mL of water per day for at least two weeks. Mouse weight was checked daily to avoid dehydration. Two days before surgery, mice got free water access. The surgery had two steps. On day 1, we implanted a head bar for head-fixation of mice. On day 2, after recovery from the head-bar surgery, we implanted the cranial window over the auditory cortex.

For the head-bar surgery (day 1), mice were implanted with a custom designed light-weight head bar. Mice were anesthetized with isoflurane (1.5% at induction, below 1% to maintain) with an additional analgesic (meloxicam 2mg/kg, subcutaneous) and eye ointment. The mice were placed in a stereotaxic apparatus. The scalp was removed above the entire cortical area. The skull was cleaned with hydrogen peroxide. The head bar was attached to the skull with metabond adhesive (parkell, S380). The craniotomy surgery (day 2) was done under isoflurane anesthesia, using the headbar to immobilize the head. Eye ointment was applied. Meloxicam (2 mg/kg, subcutaneous), enrofloxacin (5mg/kg subcutaneous) and dexamethasone (2 mg/kg subcutaneous) were administered ^64^. Enrofloxacin was also applied once per week to further prevent infection after surgery. After opening the skin, lidocaine was injected to the muscle above the auditory cortex. The muscle was removed and a craniotomy was made over the left hemisphere of auditory cortex (2.9 mm posterior and 4.2 mm lateral of the bregma) with a diameter of 3 mm, without puncturing the dura mater. A 3 mm diameter glass window (CS-3R, Warner Instruments) was mounted directly onto the dura and sealed with a mixture of krazy glue and dental acrylic powder (Lang, Jet denture repair powder/liquid). After surgery, water was given freely until the mouse recovered.

After recovery from surgery, behavioral training started. Mouse weight was carefully monitored, and additional water was given after daily training to keep the weight over 85% of the pre-restriction weight.

### Methods details

#### Behavioral training

*Behavioral setup.* The setup including part numbers and 3D print files was described before ^17^. The setup was placed inside a custom sound booth by Industrial Acoustics Company (Bronx, New York). The training was done with head-restrained mice positioned over a cylindrical treadmill running on ball bearings. The rotation of the treadmill was measured with a rotary encoder (200 P/R, Yumo). Two speakers (Avisoft Bioacoustics) were placed diagonally in front of mice for auditory stimulation. The speakers were calibrated with a free-field microphone (Type 4939, Brüel and Kjaer) ^65^. Water was delivered through 19 gauge stainless steel tubing connected to solenoid valves (Lee Company) located outside the sound box. Water was calibrated weekly and was delivered through three spouts connected to a custom lick detection circuit. The behavioral system was controlled by a custom Matlab (Mathworks) program running on Bpod framework (https://sanworks.io) in Linux.

*Task structure.* The tone frequency discrimination task required mice to select the left or right spout depending on the frequency of the sound stimulus. Mice were required to withhold licking for 0.5 sec before a trial start. A blue LED indicated the trial start (end of inter-trial-interval) and mice were required to lick the center spout to start a sound stimulus with a delay of 0.1 to 0.3 sec. The sound stimulus was a ‘tone cloud’ stimulus as described before ^66^ (see below). At the end of the tone cloud, mice received a small reward of water (0.5 μl) at the center spout. During the sound stimulus, mice were allowed to lick any spout. When the tone cloud contained more low tones than high tones (low category tone), the selection of left or right spout provided a large reward (2 μl of 2% sucrose water) (correct) or a noise burst (0.2 sec) (error), respectively. The high category tone cloud had the opposite correct and error choices. The time between the side lick and reward/noise varied between 0 and 0.2 sec. The interval of tone cloud between trials was at least 5.4 sec (6.6 sec in median), except in 1 out of 96,420 trials in 166 sessions (4.4 sec), to eliminate a sound adaptation in the auditory cortex ^28^. When mice did not select the side spout within 30 sec from the trial start, a new trial started.

*Stimulus generation.* The tone cloud was 0.6 sec long and consisted of a series of 30 ms pure tones with rise/decay ramps of 3 ms, presented at a rate of 100 tones per second. The frequency of each tone was sampled from the bottom 6 and top 6 tones of 18 logarithmically spaced slots (5 to 40 kHz). The tone cloud in each trial contained the low (5 – 10 kHz) or high frequency tones (20 – 40 kHz), and was categorized as low or high depending on the dominant frequency. The proportion of high tones in each tone cloud was selected from 6 settings (0, 0.25, 0.45, 0.55, 0.75, 1) with the probability of (25%, 12.5%, 12.5%, 12.5%, 12.5%, 25%, i.e., 2:1:1:1:1:2). In the stimulus probability task, we changed the probability between categories (see below) but kept the stimulus probability within the category constant (i.e., 2:1:1). The intensity of tone cloud was constant in each trial, but sampled from either 70, 75 or 80 dB SPL (sound pressure level in decibels with respect to 20 μPa) to discourage mice from using loudness to solve the task.

*Stimulus probability and reward amount tasks.* Every session started with an easy block where only the 100% low or high tone clouds were presented 60 to 80 trials with the stimulus probability of 50%-50% (low-high). The stimulus probability task then changed the stimulus probability of the low or high category tones in blocks of 175 to 220 trials (200 trials in 70 out of 83 sessions). The stimulus probability of one block started with either 70%-30% (low-high, left block) or 30%-70% (right block), and the probability reversed in the next block. After mice experienced the two blocks, the stimulus probability became 50%-50% for the rest of the session (post block). The reward amount for the correct choice was constant in all trials (2 μl).

The reward amount task changed the reward amount of the left and right spout in blocks, while the stimulus probability was 50%-50% (low-high) in all trials. In each block, the reward amount for the left-right correct choice was either 3μl-1μl or 1μl-3μl (left or right block). The block schedule was the same as in the stimulus probability task.

*Training schedule.* The initial phase of training was described previously ^17^. On the first day of training, we used only stimuli with 0.1 or 0.9 proportion high tones and free sucrose water from the correct side spout (free-reward trials). From the second day of training, we mixed the free-reward trials and the choice trials which required mice to select side spouts to get reward. Mice learned to lick the side spouts independently, presumably by feeling the delivery of the free-reward and licking towards it. We gradually decreased the proportion of free-reward trials. Based on performance (no strict criteria), we introduced more difficult tone clouds. The inter-trial-interval was then increased gradually by 3 days of training. We introduced the stimulus probability task and reward amount task after mice succeeded in getting reward in the task without any manipulations of stimulus or reward (about 90 % correct in 100 % low or high tone clouds). Before starting the imaging sessions, mice were fully trained on the stimulus and reward amount tasks.

*Recording schedule.* Each mouse performed both the stimulus probability task and reward amount task. We imaged the same field of view (FOV) and neurons during both tasks as follows: we imaged from one FOV during the stimulus probability task (day 1) and reward amount task (day 2), then switched to another FOV during reward amount task (day 3) and stimulus probability task (day 4). Day 5 had the same procedure as day 1 and so on. We did not use the following sessions for analysis: (i) mice did not complete the two blocks with opposite stimulus probability or reward amount in a given session, (ii) mice made errors in more than 25 % of trials with either the 100 % low or high tone cloud. In these cases, the same combination of FOV and task was selected the next day. Exceptionally, the same FOV was imaged in 11 and 2 sessions in the stimulus probability and reward amount tasks, respectively. In these sessions, we selected one session for analysis only based on the behavior data without analyzing the imaging data. Also, sessions described below were not used in the analyses: (i) the FOV was imaged only during either stimulus probability or reward amount task (7 and 1 sessions in stimulus and reward task), (ii) the FOV had a few bright cells which might indicate over-expressed GCaMP6f (1 session each task), (iii) the difference of imaging date between the two tasks was 17 days (1 session). In total, we analyzed 83 sessions in both the stimulus probability and reward amount task. The difference of imaging date between tasks was typically one day (1 day difference, 60 session pairs; 2 days, 15 pairs; 3 days, 7 pairs; 4 days, 1 pair).

#### Wide-field imaging

To identify the position of primary auditory cortex and determine the locations for two-photon imaging, we conducted one-photon wide-field calcium imaging through the chronic window. Mice were awake and head-fixed on the treadmill. Two blue LEDs were used for illumination through fiber guides directed on the window. Emitted photons were captured by a CCD camera (Vosskuehler 1300QF). Frames were acquired at 4 Hz using a custom Labview software (National Instruments). Sound stimuli were presented at approximately every 6 s. Each stimulus was a 2 s train of pure tone pulses (20 Hz) at the frequency of either 4, 11 or 32 kHz with the intensity of 70 dB SPL. The sound evoked activity (*F*) was analyzed in each pixel as follow:

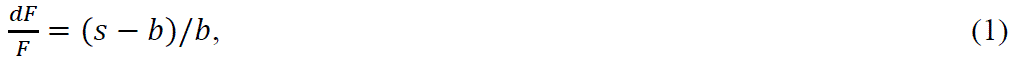

where s and b were the average intensity of stimulus (2 s, 8 frames) and pre-stimulus frames (2 s, 8 frames), respectively. Each frequency tone was repeated 10 times in a pseudo random order (30 stimuli in total). The 4-kHz pure-tone evoked a characteristic constellation of activity in primary, anterior, and secondary auditory fields (A1, AAF and A2 (or SRAF))^22,23^. The most posterior activity spot was identified as A1 (**Figure 2**).

#### Two-photon imaging

After training in the task, we started imaging experiments. As we observed almost no clearly over-expressing cells with GCaMP6f in our transgenic mice, we continued the experiments for 4 to 12 weeks. We imaged 9 to 21 fields of view (FOVs) from 2 to 3 selected XY locations per mouse (**Figure 2 and S2**). The depths of FOVs were between 110 and 510 μm. The FOVs were mainly from layers 2 and 3 (79 out of 83 sessions below 400 μm). On each experimental day, we imaged one FOV of one location.

Imaging was performed using a custom-built two-photon microscope with the objective lens at an adjustable angle to image auditory cortex without tilting the mouse. The resonant scanner was outside of the sound box, rendering it inaudible inside the box. We used a 20x/N.A. 1.0 water immersion objective for imaging (Olympus). A Ti:sapphire laser (Chameleon, Coherent) was operated at 910 nm to excite fluorescence, which was detected with GaAsP photomultiplier tube (Hamamatsu) in the spectral range of 500 – 540 nm (Chroma ET520/40m-2p). A 12kHz resonant scanning system was used to acquire images at 45 Hz at a resolution of 512 x 512 pixels corresponding to a 380 x 550 μm^2^ field of view with the 20x objective. The microscope and image acquisition were controlled by an open source software (ScanImage, Vidrio Technologies, ^67^). In each task session, we imaged for about one hour continuously in more than 500 trials.

When slow drift of the imaging plane was observed, the objective position was manually adjusted (typically 1 μm at a time) during the inter-trial interval to match the recording site as precisely as possible to an average image taken at the beginning of the session.

#### Data analysis

All analyses were conducted with Matlab (MathWorks). In the figures, error bars of the mean represent standard deviation or standard error of the mean (s.e.m.). Error bars of the median represent median absolute deviation (MAD) or robust standard error (1.4826*MAD/sqrt(n); n = number of data points) ^68^.

*Behavioral analysis.* In every behavioral task session (one per day, one imaging plane at one location), trials in which mice succeeded to select the left or right spout were analyzed. In total, we analyzed 166 sessions from 6 mice from the stimulus probability task (83 sessions) and reward amount task (83 sessions) (mouse1, 20 sessions; mouse2, 42 sessions; mouse3, 28 sessions; mouse4, 24 sessions; mouse5, 34 sessions; mouse6, 18 sessions). Each field of view (FOV) was imaged during the two tasks.

We used a logistic regression to quantify the behavior bias between blocks (psychometric function) (**Figure 1**). The same equation was used to analyze the neurometric function of single and population neurons described later (**Figure 4 and 6**) ^69^:

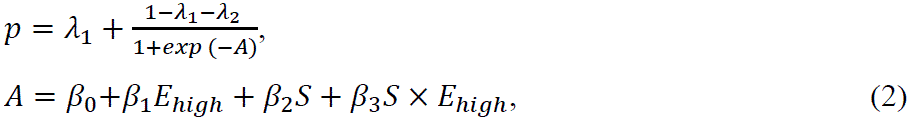

where *p* was the probability to select the right spout. β_0−3_ were regression coefficients. β_1_determined the slope of the psychometric curve of behavior (stimulus sensitivity). β_2_quantified the choice bias between the left and right blocks, while β_3_ quantified the change of stimulus sensitivity by blocks. λ_1_and λ_2_were lapse rates which were λ_1_ = λ_2_ for the model fitting in **Figure 1E**. *E_high_* was the proportion of high frequency tones in a tone cloud. *E_high_* had 6 settings (0, 0.25, 0.45, 0.55, 0.75 1). *S* was –1 or 1 for the left or right block. For model fitting, we used the trials only during left and right blocks.

To determine which parameters were relevant for mice behavior, we used a likelihood ratio test. We first averaged the log likelihood of the logistic-regression model across sessions in each mouse and then averaged across mice (**Figure 1E**) ^70^. The parameters were set to achieve the maximum likelihood. In addition, we modeled the mice behavior in each block with the full-parameter logistic regression model (equation 2). This full model was used to analyze the difference of right choice probability between blocks (Δ fraction rightward) based on the average choice probability in logistic regression in each block (**Figure 1D and 6**). As the full model was independently applied to the data in each block, β_2_ and β_3_ were set to 0 (4 parameters in total).

*Neural analysis.* We used an open-source software, Suite2P, for the motion correction and extraction of regions of interest (ROIs) from raw imaging data (https://github.com/cortex-lab/Suite2P) ^71^. The parameters for Suite2P were default except the diameter of ROIs as 15. ROIs were then manually extracted with the GUI in Suite2P. Suite2P also detected the overlapping ROIs between the images taken during the stimulus probability task and reward amount task (registers2P). The parameters for overlap detection were default (proportion of overlap, 0.6). In each ROI, a neuropil correction was done based on Suite2P. 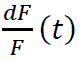 was calculated based on the signal at frame *t*, *F*(*t*), as follows:

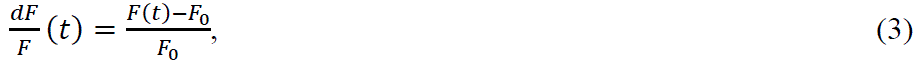

where *F_0_* was the average signal during 1 sec (45 frames) before the LED onset (trial start) in each trial.

*Task-relevant neurons.* For every ROI, 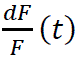 were analyzed at the following 6 time windows during the task to investigate task-relevant neurons: (i) between the LED and sound onsets; (ii) during the sound presentation (0.6 sec); (iii) between 0 and 1 sec from the choice; (iv) between 1 and 2 sec from the choice; (v) between 0 and 1 sec from the reward or noise-burst delivery; (vi) between 1 and 2 sec from the reward or noise-burst delivery. The neural activity was analyzed in the following trials in each time window: (i, ii) all, low-, or high-category-tone trials; (iii, iv) all, left-, or right-choice trials; (v, vi) all, reward, or noise-burst trials (3 conditions in each time window). This analysis was independently applied to the activity during the stimulus probability task and reward amount task (2 tasks). We defined the ROI as task-relevant when the aligned and averaged activity had a significantly positive value at any time window (6 settings), in any condition (3 settings), and in any task (2 settings) compared to the baseline activity (36 settings in total (6 x 3 x 2)) (one-sided Wilcoxon signed rank test, p < 0.036 after Bonferroni correction in each comparison). The baseline activity was defined as the activity before the LED onset with the corresponding time window in each condition. The activity during left and right blocks was used in the analyses here and hereafter.

*Neural encoding.* We investigated the activity of sound responsive neurons which had (1) significant increase of activity during sounds (time window of (ii) in previous section) compared with the activity during inter-trial interval and (2) preferred tone cloud (p < 0.01 in Kruskal-Wallis test) in both the stimulus probability and reward amount tasks. The preferred tone cloud of each neuron was defined as the stimulus with the highest average activity in correct trials. The activity of sound responsive neurons were compared between left and right blocks to identify the block (context) modulated neurons which had significant change of activity between blocks at the preferred tone cloud (p < 0.05, two-sided Mann Whitney U-test) (**Figure 3**). The activity was also compared between correct and error trials (**Figure S6**).

*Signal and noise correlation.* Signal and noise correlations were investigated using the Pearson correlation coefficient between pairs of sound-responsive neurons. Signal correlation was defined as the correlation coefficient between the mean activity of each 6 tone cloud in a given neuron pair^33^. For the noise correlation, the calcium traces for each of 6 tone clouds were independently z-scored (mean subtracted and divided by the standard deviation) to get the variability of activity in each stimulus. The noise correlation was defined as the correlation coefficient between the variability (noise) of neuron pairs.

*Single neuron decoding.* Our decoder for single neurons used a simple threshold to categorize the neural activity into low and high tones. The average activity during sound was used in the decoder. To compute the decoding performance of single neurons (**Figure 4**), the decoding threshold was computed from the training data using 10-fold cross-validation and tested in the test data. The cross validation was applied 100 times to reduce the variance of decoding performance from the random grouping. To investigate the relationship between the decoding threshold, decoding performance, and block-dependent modulation (**Figure 7**), we investigated one threshold using all the data. The block modulation was defined as the difference of median activity between blocks at the preferred tone cloud. The decoding threshold and block modulation were z-scored (mean subtracted and divided by the standard deviation) for population analysis.

*Population neural decoding.* We used a sparse logistic regression (SLR) to decode sound category (low or high) from the population activity of task-relevant neurons. We used a software package, sparse learning package (SLEP) (https://github.com/jiayuzhou) ^72^. First, a logistic regression provided the likelihood of decoding performance as follows:

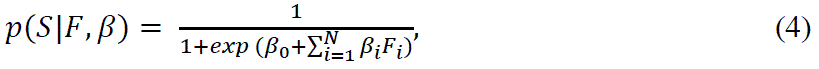

where *S* was the tone category. *F_i_* was the average activity (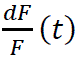) of each neuron during sounds. *N* was the number of task-relevant neurons in each session. β_*i*_ was the coefficient for neuron *i*. The SLR minimized the following equation with the regularization parameter λ:

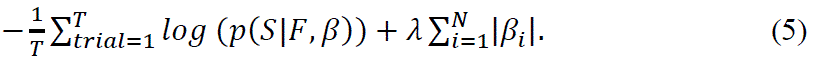

We used nested 10-fold cross validation to evaluate the decoding performance ^31^. First, trials of left and right blocks in one session were equally divided into 10 groups. The 9 groups of data were used to train the SLR, while the remaining 1 group was used to validate the decoding performance. We repeated all the 10 combinations of training and test data to evaluate the performance in all trials. The regularization parameter λ was determined with a 10-fold cross validation within the training data (9 groups of original data), such that the test data (the remaining 1 group of data) was neither used to determine λ nor β. The nested cross validation was applied 100 times to reduce the variance of decoding performance from the random grouping. The likelihood of SLR was binarized at 0.5 (decision threshold) to get the correct rate in each session. The binarized outputs were directly used as the choice outputs in the no-context model, while the non-autonomous choice model had the optimal block-dependent decision thresholds for choices (**Figure 6E**). The neurometric function in each block was analyzed based on the binarized likelihood in SLR. Choice bias in the neurometric function was analyzed with the full-parameter logistic regression model which was independently applied in each block (equation 2) (**Figure 4 and 6**).

Same as the sound decoding, the choice and context decoding were performed with the nested 10-fold cross validation to estimate the choice and context in each trial, respectively (**Figure 6 and S5**). The output of context decoding in each trial was used for the context input of the autonomous choice model (**Figure S5D**). In all the sound-, choice-and context-decoding with SLR, we used the activity of task-relevant neurons.

We investigated the sound decoding performance on shuffled neural data to assess the effect of noise correlations on population neural decoding (**Figure 5D**). In every task-relevant neuron, we swapped the activity for 2 trials with the same tone cloud and investigated whether the swap increased or decreased the correlations among neurons. If the average population correlations decreased with the swap, we accepted the swap and otherwise rejected. In each neuron in each tone cloud, we repeated the swap 100 times. This shuffling changed the noise correlation among neurons but did not change the signal correlation and mean tone-evoked activity ^33^.

To compare the decoding performance of constant weights, dynamic weights, and discordant weights (**Figure 6A**), we trained the SLR by using an equal number of low– and high-category-tone trials in each block by subsampling the training data. This prevented the decoder from using the knowledge of stimulus probability for classification especially in the stimulus probability task. For training the SLR with dynamic weights, which had independent β and λ in each block, the nested cross validation was separately applied to the trials in left and right blocks. For training the SLR with constant weights, the nested 10-fold cross validation was applied once in the two blocks. The number of trials for training was adjusted such that the equal number of trials were used to train the SLR with dynamic and constant weights. Training of SLR with discordant weights was performed in the same way as the dynamic weights, but trained and tested with the different blocks.

The performance of SLR in sound decoding was compared with that of support vector machine (SVM) (MATLAB, fitcsvm), standard logistic regression (SLEP with λ close to 0), and generalized linear model (GLM) during sound (**Figure S4A**). The decoders had constant weights and used the activity of task-relevant neurons. For GLM, we used a log-linear model in which the log scale of neural activity was fit to a linear regression with task parameters (sound category, choice, block, outcome, licking frequency for three spouts (left, center, right), and locomotion speed). The relevant parameters were investigated with Lasso (software package, SLEP) with the nested 10-fold cross validation. GLM decoder then used Bayes’ theorem to decode sound category from the log-linear model, assuming that the activity of each neuron was independent ^26^.

*Sound and choice decoding during the entire task.* We investigated the decoding performance of sound and choice in 60 different time windows on trials that were selected to decorrelate the two variables (**Figure S6A-C**). Each time window had 0.27 sec (12 frames) with a time step of 0.067 sec (3 frames). To decode sound category without the choice effect, we sub-selected either left-or right-choice trials and independently decoded the sound category. To decode choice without the sound effect, we sub-selected trials with each of 6 tone clouds and decoded the choice. The decoding performance was defined as the average of the 2 and 6 cases in the sound and choice decoding, respectively. All the decoding was done with nested 10-fold cross validation in SLR. The training data was sub-sampled such that the number of trials for each category was equal. The 10-fold cross validation was applied twice in this time-window analysis. The number of training data was different between the sound and choice decoding, making it difficult to directly compare the performance of the two decoders.

#### Pure tone response

After each two-photon imaging session during the task, we investigated the pure tone responsiveness of neurons. We presented pure tones (5 to 40 kHz with 18 logarithmically spaced slots, 30 ms with 3 ms rise/decay ramps, 50 or 75 dB SPL), white noise (30 ms with 3 ms rise/decay ramps, 50 or 75 dB SPL) and the tone clouds used during the task. The tone clouds were selected to contain 10 stimuli from each of 6 tone-cloud settings. We presented each stimulus in pseudo-random order with the inter-stimulus interval of 2 s. In total, we presented 440 stimuli (360 pure tones (18 frequencies x 2 intensities x 10 times); 20 white noise (2 intensities x 10 times); 60 tone clouds (6 settings x 10 times)).

Pure tones with 75 dB SPL were used to analyze the tone evoked responses of auditory cortical neurons. Based on the fluorescence intensity *F*(*t*), 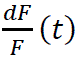 in each ROI was calculated by the same procedure as for the task (equation 3), except that *F_0_* was the average fluorescence intensity during 1 s (45 frames) before the sound onset in each trial. Neural activity following the tone presentation of 2 adjacent frequencies was analyzed together (9 frequency categories). When the activity within 1 s from the tone onset was higher than the activity before sound at least in one of the 9 frequencies, we defined the neuron as tone responsive (one-sided Wilcoxon signed rank test, p < 0.005). The best frequency (BF) of the neuron was defined as the tone frequency which evoked the highest activity.

Based on the BFs, we investigated the sound response map in the auditory cortex. In each mouse, the imaging planes of all depths were superimposed to average the BFs in XY locations (each location 200 x 200 μm with 100 μm step) (**Figure 2 and S2**).

#### Supplemental note

##### Two-photon microscopy in the auditory cortex during two tasks

We imaged calcium activity from six mice expressing GCaMP6f in excitatory neurons. All the mice performed both tasks. Because the auditory cortex is located on the side of the head, the objective lens for imaging was placed diagonally, allowing the mouse to remain in a more comfortable configuration parallel to the ground (**Figure 2A**). We first identified the location of primary auditory cortex using one-photon wide-field imaging (**Figure 2B**). A 4-kHz pure-tone evoked a characteristic constellation of activity in primary, anterior and secondary auditory fields (A1, AAF and A2 (or SRAF)) ^22,23^. The most posterior activity spot was identified as A1, which was target of further detailed study using two photon microscopy.

In each field of view (FOV), we investigated the frequency tuning of neurons by presenting pure tones with various frequencies (**Figure 2C**) to passive animals. We imaged three xy planes at varying depths along the z-axis to sample layers 2 and 3. On average, 13% (36 / 280) of all neurons per FOV showed tone-evoked activity in at least one frequency (p < 0.005 in one-sided Wilcoxon signed rank test); the relatively low fraction of tone-responsive neurons is consistent with previous reports in various preparations ^24–26^. We defined a neuron’s best frequency (BF) as the frequency which elicited the highest activity. An example of a BF sound response map, constructed as the average of neurons over the z-axis, is shown in **Figure 2D** (all mice, **Figure S2A**).

To image the activity of the same neurons during both the stimulus probability and reward amount tasks, we switched the task on alternate days, keeping the same FOV (**Figure 2E**). We used a software package, Suite2P with registers 2P ^71^, to detect overlapping regions of interest (ROIs) between the tasks, as well as to detect ROIs from raw imaging data. In total, Suite2P detected 23088 and 23350 ROIs in the stimulus probability and reward amount tasks, respectively (278 and 281 ROIs on average per FOV). Suite2P extracted 17523 overlapping ROIs (211 ROIs per FOV).

We identified task-relevant neurons from the overlapping ROIs (**Figure 2F**). We defined a “task-relevant neuron” as any neuron showing increased activity in at least one of the following 6 time windows compared to the inter-trial interval in either the stimulus probability or reward amount task (2 task settings) in 3 different trial types (36 settings in total (6 x 2 x 3), one-sided Wilcoxon signed rank test, p < 0.036 after Bonferroni correction). The 6 time windows were (i) before the sound onset; (ii) during the sound presentation; (iii) between 0 and 1 sec from the choice; (iv) between 1 and 2 sec from the choice; (v) between 0 and 1 sec from the reward or noise-burst delivery; (vi) between 1 and 2 sec from the reward or noise-burst delivery. The 3 different trial types in each time window depended on the sound category (i, ii: all, low, high), choice (iii, iv: all, left, right), or outcome (v, vi: all, correct, incorrect). We identified 13581 task-relevant neurons out of 17523 overlapping neurons (78 % of ROIs per FOV on average). The peak activity time and signal strength of task-relevant neurons were correlated across days (Spearman partial correlation eliminating the effect of mice or sessions, time: r = 0.56, p < 1E-10, strength: r = 0.71, p < 1E-10) (**Figure 2G and S2B**), consistent with a previous finding in mouse parietal cortex ^27^. When we focused on the time window at sound onset, the proportion of neurons that significantly increased the activity in both the tasks was 27.5 % on average which was larger than that during passive sound presentations after both tasks of 6.6 % (p = 3.6E-15 in linear mixed-effects model, 6 mice, 83 sessions).

##### Sound responses are modulated by the stimulus and reward context

We next tested if activity in auditory cortex was modulated by the block-wise changes in stimulus or reward context. We set the median inter-sound-interval to 6.6s to minimize changes in sound-evoked activity due to task-unrelated sound adaptation^28^. Among the task-relevant neurons, we focused on sound-responsive neurons that showed increased activity during stimulus presentation in the stimulus probability and reward amount tasks compared to the activity during inter-trial interval, and that also had significantly different activity in at least one tone cloud compared to the rest (p < 0.01 in Kruskal-Wallis test) (one session in **Figure 3A**, 83 sessions in 2909 and 2573 neurons; 17% and 15% per FOV on average). Of these, 1735 neurons (11% per FOV) showed sound responses during both tasks. For each sound-responsive neuron, we defined the “preferred tone cloud” as that stimulus which elicited the highest activity among all the tone clouds. The preferred tone cloud was preserved between the two tasks (**Figure S2C,** Spearman partial correlation, r = 0.81, p < 1E-10), and correlated with the best frequency during passive pure tone presentation (**Figure S2D**, r = 0.32, p = 1.4E-14 in stimulus task; r = 0.36, p = 1.4E-17 in reward task). The fraction of neurons responsive to sounds during the task was comparable to the fraction responsive to pure tones during passive listening (13%) but much smaller than the 78% of neurons that were modulated by the task (task-relevant neurons). Thus, consistent with previous findings in auditory cortex ^25^, most task-responsive neurons were not sound-responsive.

We investigated the contextual modulation of sound responses by comparing the activity between left and right blocks within each task. A representative sound-responsive neuron showing increased activity in the right block compared to the left block during the stimulus probability task is shown in **Figure 3B**. We found that the activity of 35.7% of sound responsive neurons was modulated by the changes in stimulus-probability or reward-amount (**Figure 3C**). Context modulation of sound responsive neurons was greater than sound-unresponsive task-relevant neurons (0.089 +/− 0.16 vs 0.025 +/− 0.030, stimulus modulation; 0.048 +/− 0.075 vs 0.024 +/− 0.027, reward modulation; mean +/− std) (**Figure 3D**). We then aligned the activity of each sound responsive neuron based on its “preferred block,” i.e. the block associated with the preferred tone cloud stimulus for that neuron. Specifically, the preferred block was “left” for neurons whose preferred tone cloud was in the “low” category and “right” for neurons whose preferred tone cloud was in the “high” category. We categorized the neurons by their preferred tone difficulty and compared activity between blocks elicited during the preferred tone cloud stimulus. We found that the activity could be modulated in either the positive or negative direction (**Figure 3E**). Contextual modulation was stronger in neurons tuned to the difficult or moderate tone clouds and weaker for those tuned to easy tone clouds (**Figure S3A,B**), irrespective of choices (**Figure S3C-E**). We then analyzed the median block modulations of sound-responsive neurons in each session (**Figure 3F**). This population block modulation was larger in the reward task than in the stimulus task. These results indicate that the encoding of sound by neurons in auditory cortex is modulated by stimulus or reward expectation and that the magnitude of the neuronal modulation is correlated with the behavioral bias (**Figure 1D**).

## Quantification and statistical analysis

All the statistical tests in this study were performed in Matlab 2016b. We mainly used two-sided statistical tests; cases where we used a one-sided test are clearly noted in the text. In tests with multiple comparisons, we set the significance threshold according to the Bonferroni correction. Statistical tests in Matlab generated p = 0 when there were small p values. In these cases, we described the p values as p < 1E-10. In the statistical tests with behavioral sessions across 6 mice, we used a linear mixed-effects model to take into account the effects of sessions imaged from the same mice as the random effects^18^. In the statistical tests with neurons across sessions and mice, we used a linear mixed-effects model to consider the random effects of sessions and mice. Data collection and analyses were not performed blind to the conditions of the experiments. No statistical methods were used to pre-determine sample sizes but our sample sizes are similar to those generally used in the field.

## Additional Resources

Not applicable.

