## Supplemental Figure1 – 7 for "Stable sound decoding despite modulated sound representation in the auditory cortex"

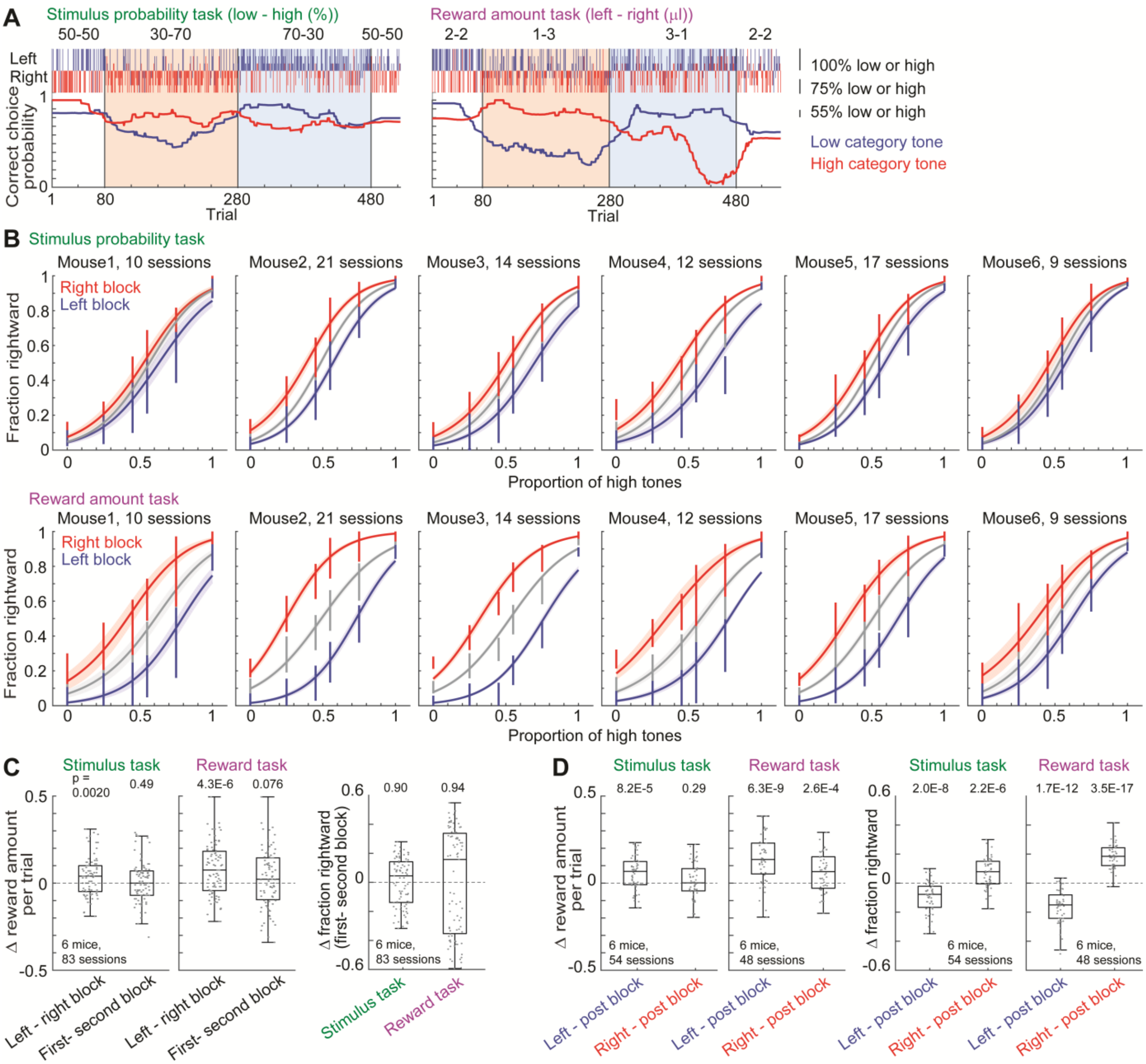


**Figure S1. Choice behavior in each mouse, Related to Figure 1**

(**A**) Block structure and choice behavior in an example session. Top panels show the choice, tone category, and tone difficulty in each trial. Bottom panels show the moving average of correct choice probability for low and high category tones. (**B**) Means and standard errors of psychometric functions are shown in each mouse. Logistic regression analyzed the psychometric function in each session ('Choice' in **Figure 1E**). Error bars show the standard deviations of choice probability per session. (**C**) Comparison of choice performance between the left and right blocks or between the first and second blocks. Choice performance was better in the left than in right blocks, while the order of the blocks did not significantly affect the performance (left: linear mixed-effects model). The order of blocks did not significantly affect the choice biases (right: linear mixed-effects model). (**D**) Comparison of choice performance between the asymmetric blocks and the post blocks which had the equal stimulus probabilities (50% - 50%) and reward amounts (2μl - 2μl). After mice experienced both the left and right blocks, we presented the post block in each session. Sessions with more than 100 trials in post blocks were included in the analyses. The choice performance in the left and right blocks were better than that in the post blocks. The choices in the left and right blocks were biased to the left and right sides, respectively, compared with the choices in the post blocks (linear mixed-effects model).

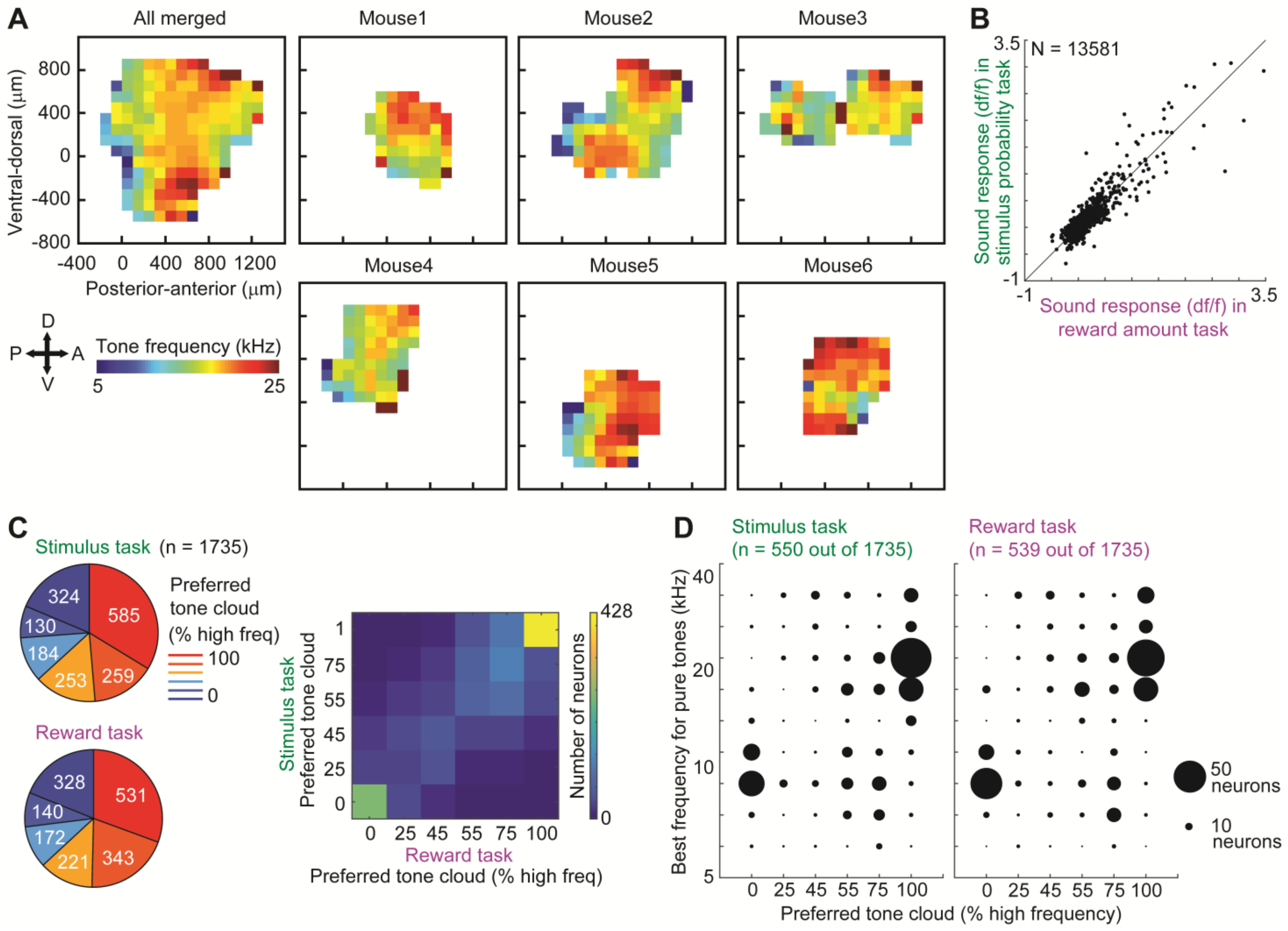


**Figure S2. Sound responses of auditory cortical neurons during passive sound presentations and task, Related to Figure 2 and 3**

**(A)** Sound response map in each mouse. In each mouse, the best frequencies of pure-tone responsive neurons were superimposed in depth to make the sound response map in XY plane. Based on the 4-kHz and 70 dB SPL tone evoked activity spot in the primary auditory cortex, the sound response map of each mouse was superimposed to make the merged map. This merged map was not used in the further analyses, as the maps across mice were not precisely aligned in the coordinates.
**(B)** Fluctuation of signal strength of neurons between stimulus probability and reward amount tasks. Median signal strength of single neurons during sounds (0.6 s) in all trials was compared between the tasks (Spearman partial correlation eliminating the effect of mice or sessions, $r = 0.71$, $p < 1E-10$).
**(C)** Preferred tone cloud of neurons that were sound responsive during both the stimulus and reward tasks ($n = 1735$). Colormap showed the preserved preferred tone cloud across tasks (Spearman partial correlation, $r = 0.81$, $p < 1E-10$).
**(D)** In the stimulus probability and reward amount tasks, 550 and 539 out of 1735 sound responsive neurons showed significant sound responses during passive listening outside the task. The pure-tone best frequency (BF) was defined as the frequency with the largest activity. When the neurons preferred the 100% high- or low-frequency tone cloud, they tended to have the high or low BFs, respectively (Spearman partial correlation, $r = 0.32$, $p = 1.4E-14$ in stimulus task; $r = 0.36$, $p = 1.4E-17$ in reward task). The tone clouds in our task had frequencies between 5-10 kHz and 20-40 kHz, with no tones between 10 and 20 kHz. Few neurons had a best frequency between 10 and 20 kHz, which might be due to the extensive training in the task.

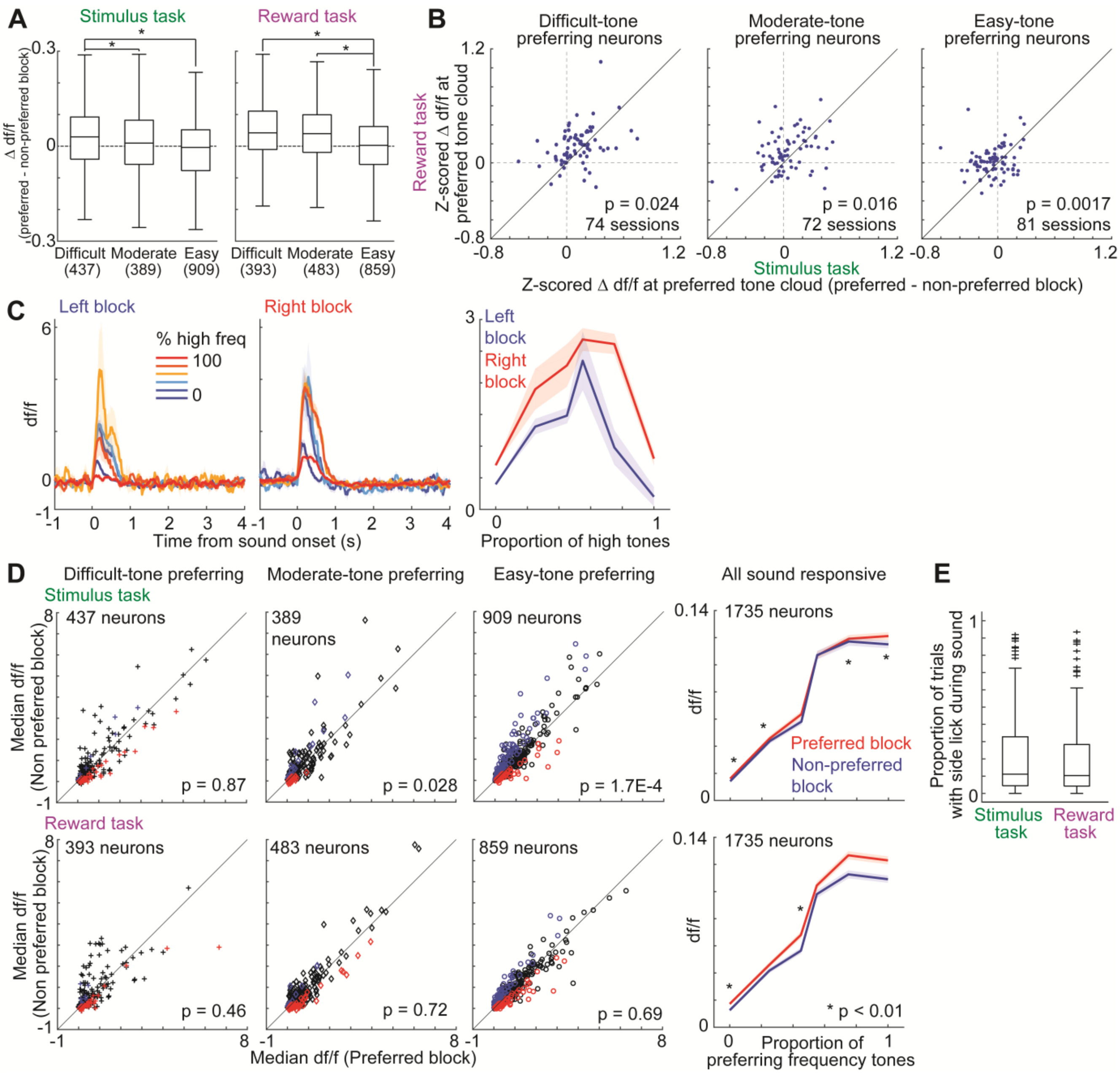


**Figure S3. Contextual modulation of sound responsive neurons, Related to Figure 3**

(**A**) Contextual modulation depends on the preferred tone cloud of neurons. We compared the contextual modulation of sound responsive neurons in different preferred tone clouds. Sound responsive neurons were categorized to the (i) difficult, (ii) moderate and (iii) easy neurons depending on the preferred tone clouds in (i) 45 or 55 % high tones, (ii) 25 or 75 %, and (iii) 0 or 100 %, respectively. Difficult neurons showed larger modulation than easy neurons (central mark in box: median, edge of box: 25th and 75th percentiles, whiskers: most extreme data points not considered outliers (beyond 1.5 times the inter-quartile range), here and hereafter. Plots without the outliers) (*, $p < 0.01$ in linear mixed-effects model in 6 mice, 83 sessions). (**B**) Comparison of median block-modulated activity per session between the stimulus and reward tasks. Data presentations comply with **Figure 3F** except that neurons were categorized to the preferred tone clouds (linear mixed-effects model in 6 mice). (**C**) Traces of neuron in **Figure 3B** in correct trials. (**D**) Contextual modulation of sound responsive neurons in correct trials. Data presentations comply with **Figure 3**. (left three panels: linear mixed-effects model in 6 mice, 83 sessions; right panel: *, $p < 0.01$ in linear mixed-effects model). (**E**) Proportion of trials with side lick during sound (83 sessions in each task).

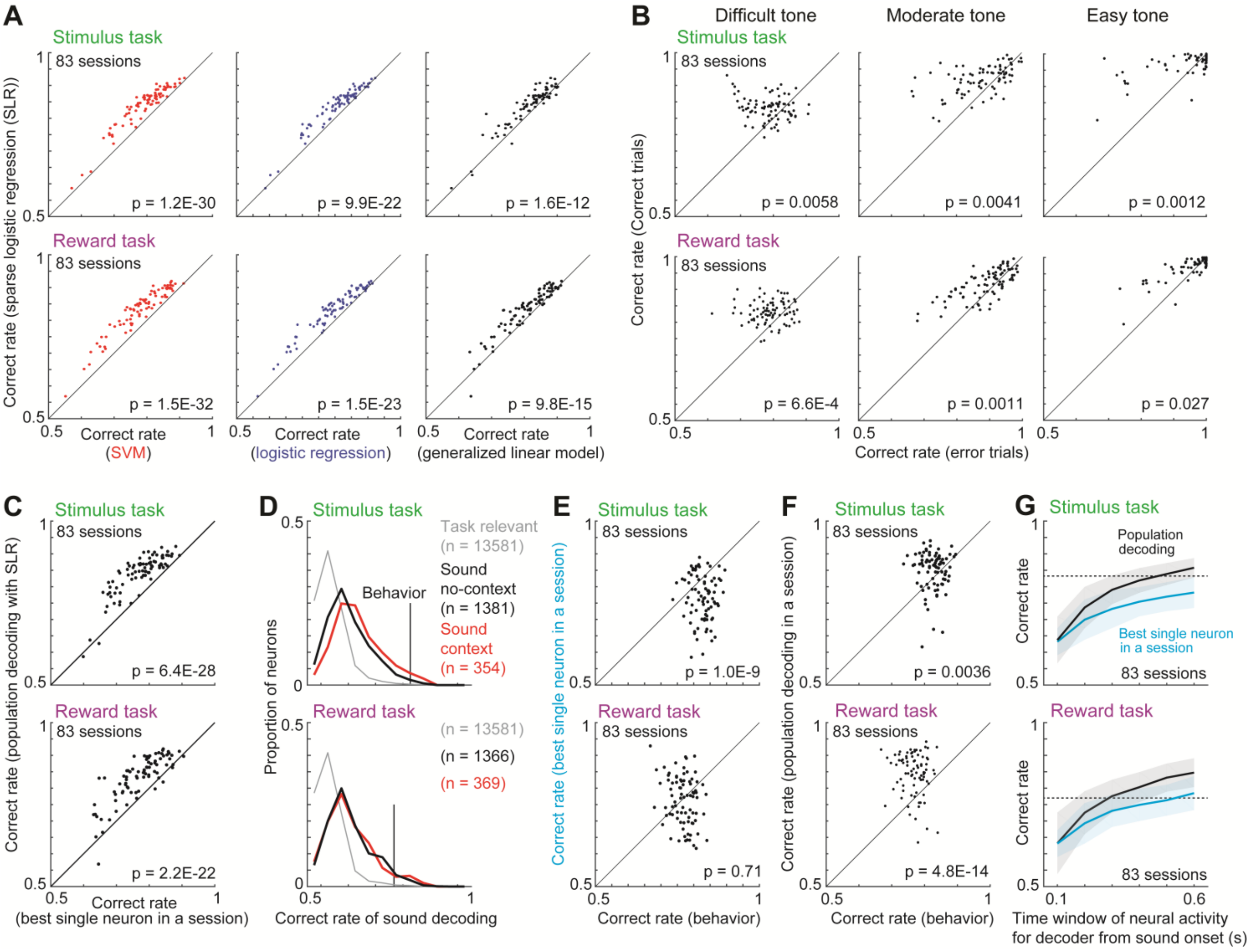


**Figure S4. Sound decoding with sparse logistic regression (SLR), Related to Figure 4**

(**A**) Sound decoding performance of SLR was compared with support vector machine (SVM), standard logistic regression, and generalized linear model (GLM). 10-fold cross validation was used (linear mixed-effects model in 6 mice, 83 sessions). (**B**) Comparison of average decoding performance of SLR between the correct and error trials (linear mixed-effects model in 6 mice, 83 sessions). (**C**) Comparison of decoding performance between population (SLR) and single neuron in each session (linear mixed-effects model in 6 mice, 83 sessions). (**D-F**) Sound decoding with various time windows. Decoders used the activity during 0.1 s at sound end. Data presentations comply with **Figure 4**. Although decoding only the final 0.1 seconds of sound yielded performance comparable to decoding the entire period (**Figure 4**), the results were confounded by the long time constant of the calcium signal, which effectively acted as an integrator. (**G**) Sound decoding performance with different length of time windows. Dot line shows the median performance of mouse behavior. The population sound decoding required the time window length of 500 ms and 300 ms in the stimulus and reward tasks to exceed the behavior. Medians and median absolute deviations.

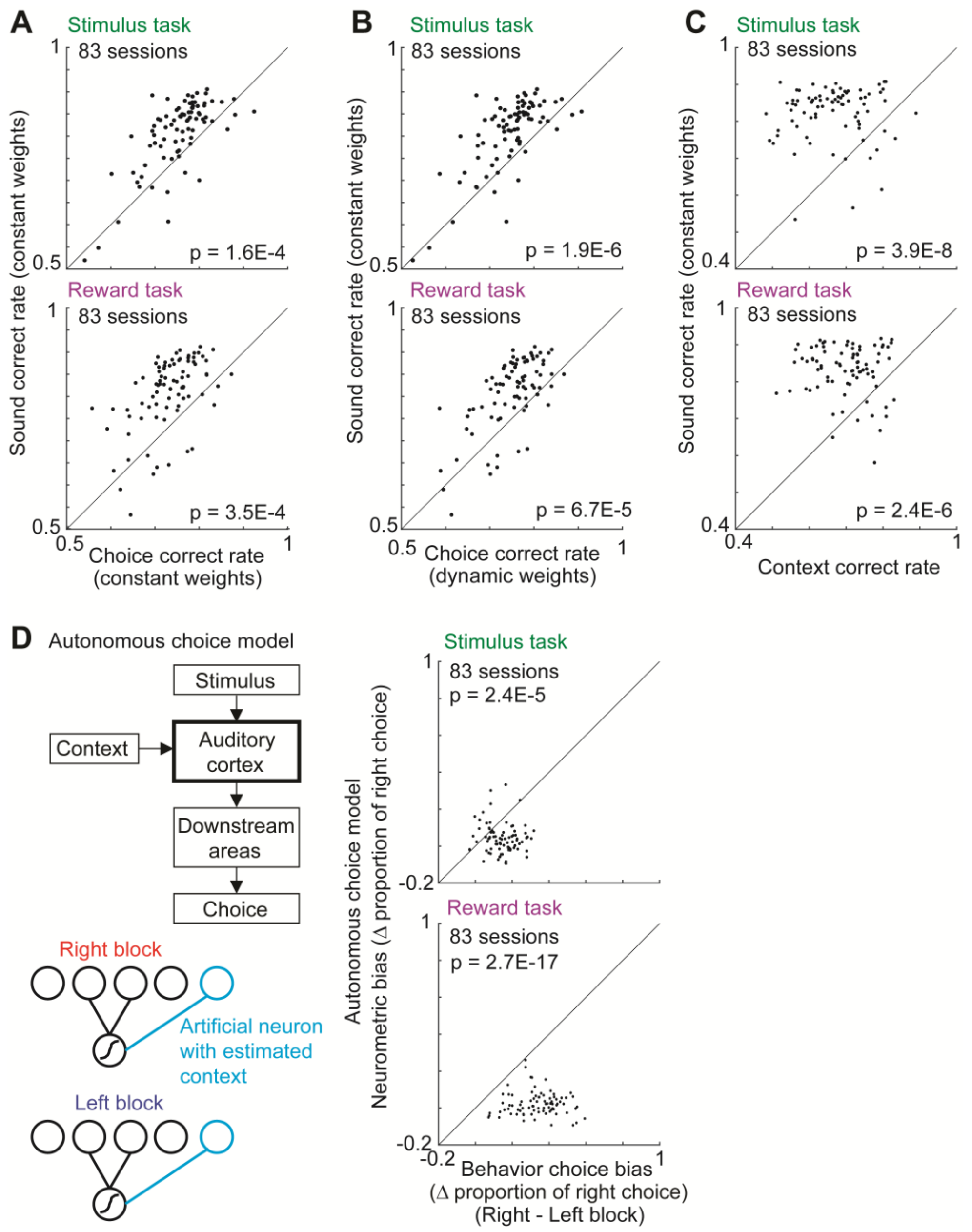


**Figure S5. Stable sound decoding from the auditory cortex, Related to Figure 6**

(**A, B**) Choice decoding with sparse logistic regression (SLR). Correct rate of sound decoding was higher than choice decoding with constant weights (**A**) and with dynamic weights (**B**) during sound (linear mixed-effects model in 6 mice, 83 sessions). (**C**) Comparison of performance in sound and context decoding (linear mixed-effects model in 6 mice, 83 sessions). (**D**) Autonomous choice model. Context was estimated from the activity of auditory cortex (cyan neuron) (**Figure 6G**) and used for the decoder. Data presentations comply with **Figure 6** (linear mixed-effects model in 6 mice, 83 sessions).

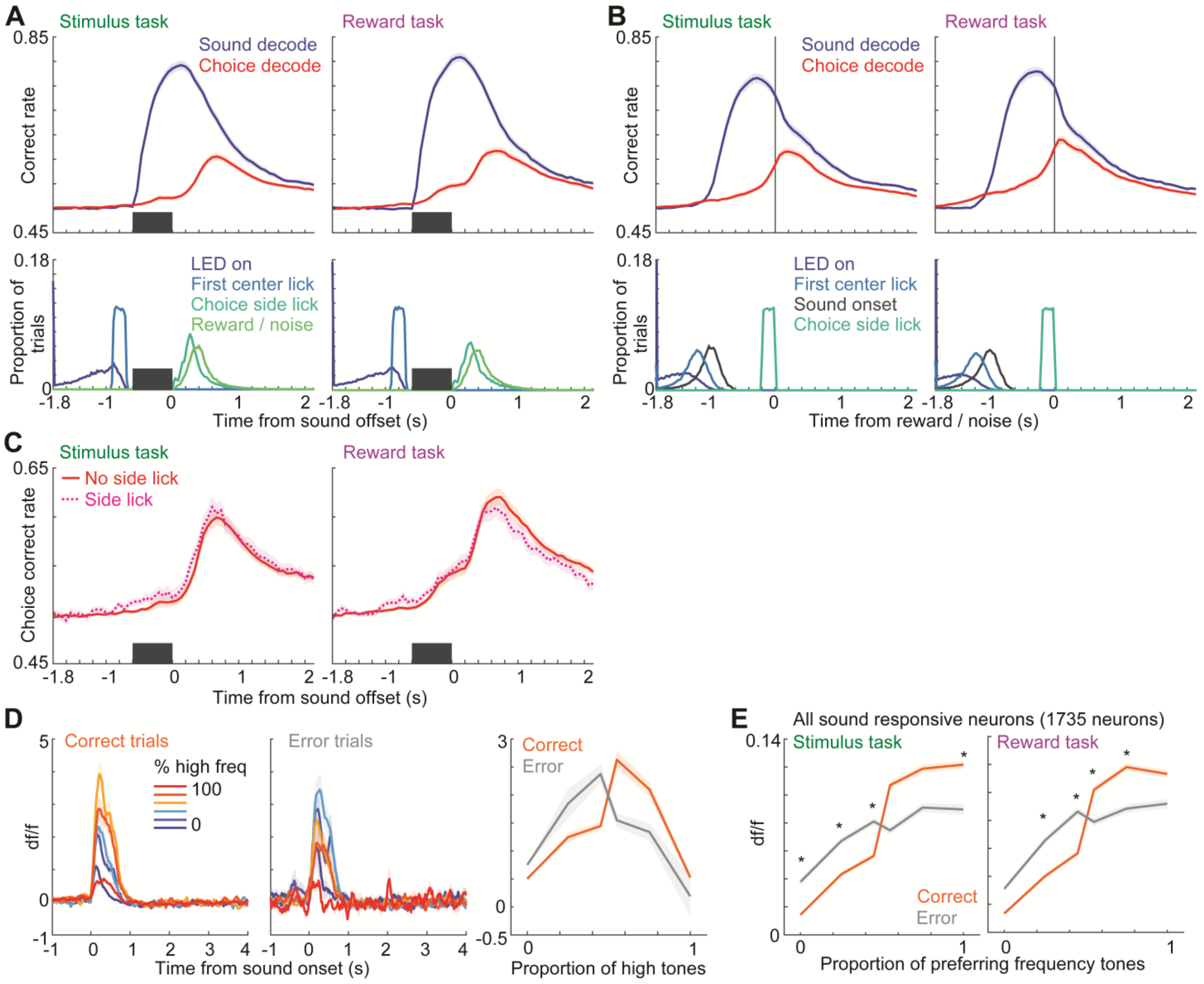


**Figure S6. Sound and choice decoding in the auditory cortex, Related to Figure 3, 6, and S5**

(**A, B**) Time course of sound and choice decoding. Means and standard errors of decoding performance are shown in different times (83 sessions in each task). Bottom panels show the timing of task variables. The decoding performance was aligned at the sound offset (**A**) and the reward / noise delivery (**B**). The performance of sound decoding reached its maximum around the sound offset, while the performance of choice decoding reached its maximum after the reward or noise burst delivery. (**C**) Choice decoding without early side licks. Correct rate of choice decoding in each session was separately analyzed for the trials with early side licks and no-side licks during sounds. Means and standard errors (83 sessions in each task). Mice licked the side spouts already during sound on some trials (**Figure S3E**), but similar choice decoding was observed in trials without early side licks.

(**D**) Choice modulation of neuron in **Figure 3B**. Data presentations comply with **Figure 3** but for correct and error trials. **(E)** Choice modulation of sound responsive neurons. Data presentations comply with **Figure 3E and S3**. Medians and robust standard errors (*, $p < 0.01$ in linear mixed-effects model in 6 mice, 83 sessions).

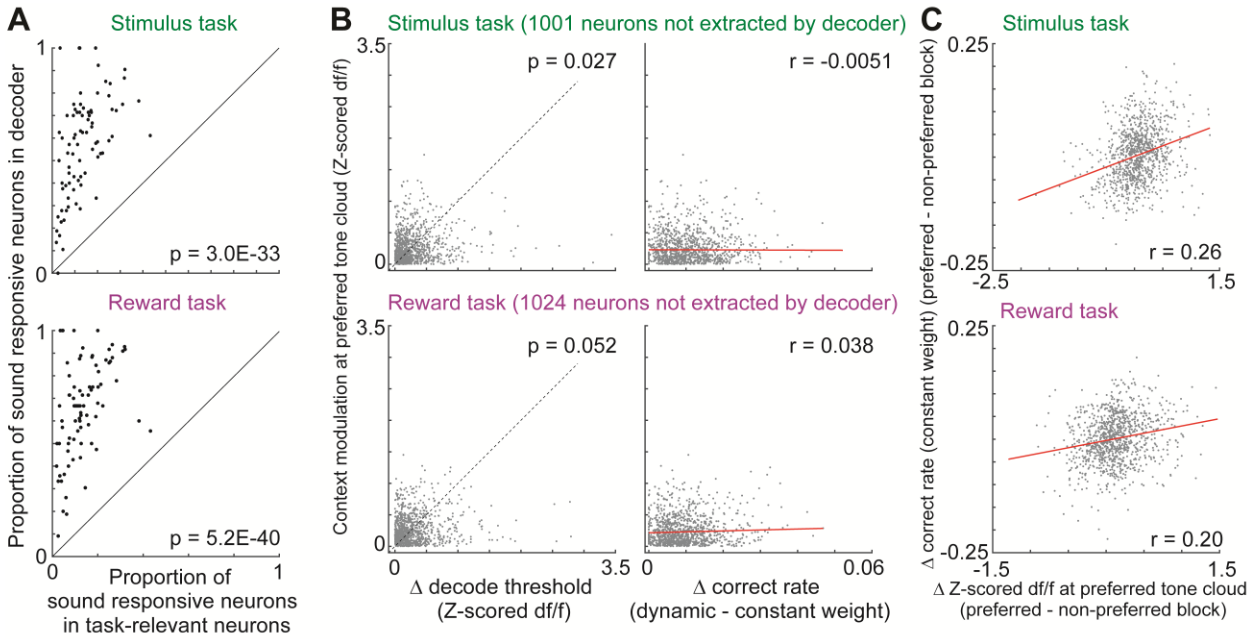


**Figure S7. Decoding threshold, contextual modulation, and correct rate of sound responsive neurons, Related to Figure 7**

(**A**) Proportion of sound responsive neurons extracted in sparse logistic regression. Proportion of sound responsive neurons was compared between the neurons extracted in the decoder and that in task-relevant neurons (linear mixed-effects model in 6 mice, 83 sessions). (**B, C**) Decoding threshold, contextual modulation, and correct rate of sound responsive neurons not extracted in the decoder. Data presentations comply with **Figure 7C,D** but for the neurons not extracted in the decoder. The contextual modulations and the changes in decoding threshold were similar (linear mixed-effects model in 6 mice, 83 sessions) (**B**, left). Spearman partial correlation between the decoding performance and contextual modulation (p = 0.87 and 0.23) (**B**, right). Contextual modulation improved decoding performance (Spearman partial correlation, p = 2.3E-16 and 6.8E-11) (**C**).
